# Visualization of individual cell division history in complex tissues

**DOI:** 10.1101/2020.08.26.266171

**Authors:** Annina Denoth-Lippuner, Baptiste N. Jaeger, Tong Liang, Stefanie E. Chie, Lars N. Royall, Merit Kruse, Benjamin D. Simons, Sebastian Jessberger

## Abstract

The division potential of individual stem cells and the molecular consequences of successive rounds of proliferation remain largely unknown. We developed an inducible cell division counter (iCOUNT) that reports cell division events in human and mouse tissues *in vitro* and *in vivo*. Analysing cell division histories of neural stem/progenitor cells (NSPCs) in the developing and adult brain, we show that iCOUNT allows for novel insights into stem cell behaviour. Further, we used single cell RNA-sequencing (scRNA-seq) of iCOUNT-labelled NSPCs and their progenies from the developing mouse cortex and forebrain-regionalized human organoids to identify molecular pathways that are commonly regulated between mouse and human cells, depending on individual cell division histories. Thus, we developed a novel tool to characterize the molecular consequences of repeated cell divisions of stem cells that allows an analysis of the cellular principles underlying tissue formation, homeostasis, and repair.

**Highlights:**

- iCOUNT reports previous cell divisions in mouse and human cells *in vitro*
- iCOUNT detects cell division biographies in complex mouse tissues *in vivo*
- iCOUNT allows for the analysis of human neural stem/progenitor cells in human forebrain organoids
- Single cell RNA-sequencing of iCOUNT cells derived from the mouse developing cortex and human forebrain organoids identifies molecular consequences of previous rounds of cell divisions

**Graphical abstract:** 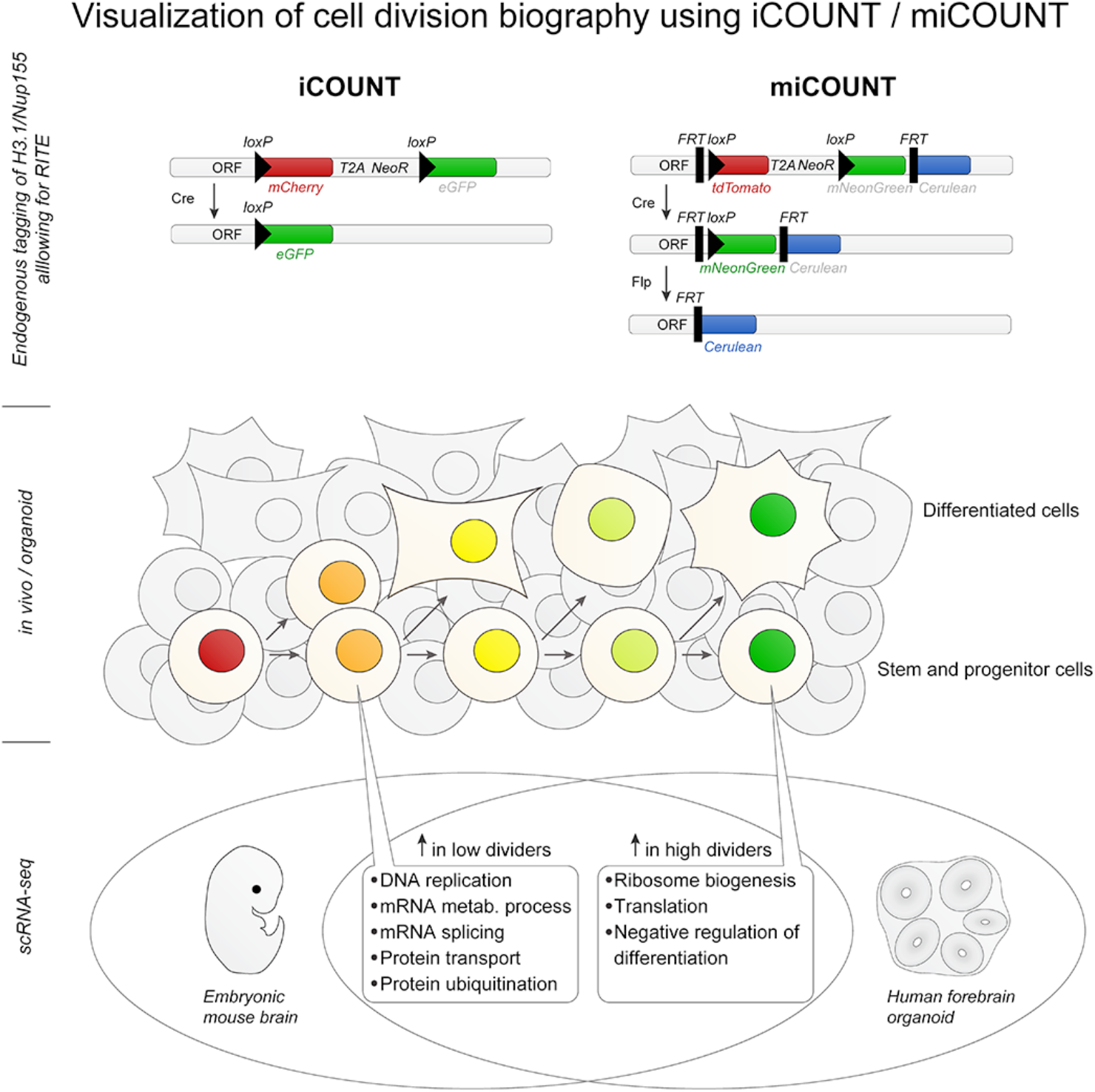

## Introduction

Somatic stem cell proliferation does not end with embryogenesis as many tissues, such as skin, intestines, the blood system, and the central nervous system continue to rely on somatic stem cells for tissue homeostasis and repair (Barker, 2014; Bianconi et al., 2013; Brack and Rando, 2012; Donati and Watt, 2015; Gage and Temple, 2013; Morrison and Spradling, 2008). Despite increasing knowledge about lineage relationships of somatic stem cells, based on advances in cellular barcoding and imaging (Fuentealba et al., 2015; Kalhor et al., 2018; Mayer et al., 2015; McKenna et al., 2016; Park et al., 2016), the functional and molecular consequences of previous cellular experiences, such as cell division events, remain largely unknown. Clearly, biographical events may be important to govern the behaviour and response to external stimuli of individual cells. Therefore, a number of tools have been recently developed with the aim to record single cell biographies based on a variety of potential experiences, for example the previous activity of multiple signalling pathways or even complete transcriptional profiles (Farzadfard and Lu, 2018; Frieda et al., 2017; Schmidt et al., 2018). This has been successful in cultured cells and bacteria using genetic approaches that allowed turning back time and to look into the past of individual cells (Frieda et al., 2017; Schmidt et al., 2018; Tang and Liu, 2018). However, a transfer of those or related technologies into more complex tissues (e.g., organoids) or even to the *in vivo* situation in mammals is missing. Previous rounds of cell divisions represent a key cellular experience of individual stem cells during organ development, tissue homeostasis, and stem cell-based cellular repair. However, how single cells respond to physiological or disease-associated stimuli, and how this response depends on an individual cell’s division history remains largely unknown. We here developed a novel genetic tool, the iCOUNT, allowing for precise measuring of previous cell division events in complex mouse tissues and human organoids on a single cell level.

## Results

### iCOUNT reports cell division events

To identify the cell division biography of individual cells in complex tissues we generated an inducible cell division counter (iCOUNT). The iCOUNT approach uses recombination induced tag exchange (RITE) of the endogenously tagged histone variant H3.1, a stable and cell cycle-dependent protein, allowing for a Cre-dependent switch from a red (H3.1-mCherry) to a green fluorescent-tagged histone (H3.1-GFP; Figure 1A) (Toyama et al., 2013; Verzijlbergen et al., 2010). We hypothesized that, after addition of Cre recombinase, every subsequent cell division reduces the amount of pre-existing red histones by one half and refills the pool of histones with newly synthesized, green histones, thus inferring the number of previous cell divisions by changes in red/green ratios (Figure 1B).

**Figure 1:**
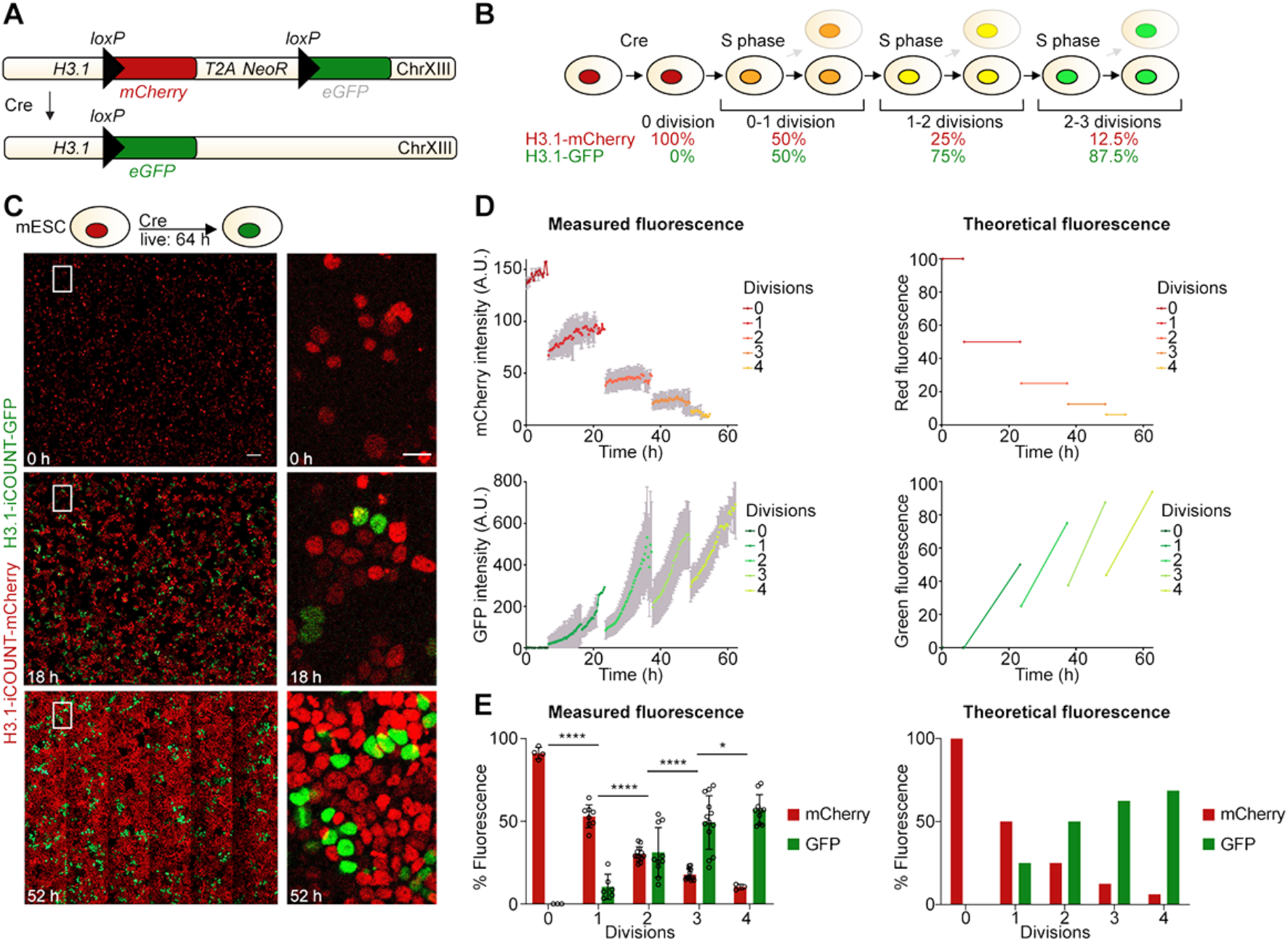
iCOUNT reports cell division events. (A) iCOUNT genetic knock-in design. (B) Theoretical change in fluorescent ratios with subsequent divisions in iCOUNT-targeted cells. (C) Selected time points of live imaging of mouse ESCs for 64h. Right panels show high magnifications of boxed areas. (D) Measured (left) and theoretical (right) changes in fluorescence intensities for red (mCherry) and green (GFP) histones of single cells (mean ± SD). (E) Quantification of measured (left) and theoretical (right) changes in percentage of fluorescence for red and green histones of iCOUNT-targeted cells (mean ± SD; open circles depict values of single cells). \**p* < 0.05; \*\*\*\**p* < 0.0001; Scale bars represent 100μm (C, left panels) and 20μm (C, right panels).

We first tested the iCOUNT system in mouse embryonic stem cells (mESCs). mESCs with H3.1-tagged iCOUNT expressed red histones and switched to green histones 24h after Cre expression (Figure S1A). Fluorescent time-lapse imaging of iCOUNT expressing mESCs over a period of 64h (Figure 1C and Movie S1) revealed that the red fluorescence dropped by half on every cell division and was refilled by green fluorescence (Figure 1D-E, left panels), confirming the theoretically expected values (Figure 1D-E, right panels). Tagged histones were symmetrically segregated between daughter cells (Figure S1B). Comparing the directly-measured division history of individual cells with the analysis of the percentage of green histones at each time point showed that the iCOUNT system correctly predicted 95.5% of all division events (Figure S1C). Using fluorescence-activated cell sorting (FACS), we analysed long-term dynamics (6 days post-Cre) of the colour exchange at the population level and found a robust shift from red to green fluorescence in cells exposed to Cre recombinase (Figure S1D-E).

To test the versatility of RITE-based cell division counting, we tagged NUP155, a stable core component of the nuclear pore complex, with the iCOUNT cassette (Toyama et al., 2013). NUP155-iCOUNT showed a similar colour exchange as H3.1 after addition of Cre, as measured by time-lapse imaging and FACS analysis in mESCs (Figure 2A-C). To test the ability of iCOUNT to report cell divisions in other cell types, we generated H3.1-iCOUNT and NUP155-iCOUNT mouse hippocampal neural stem/progenitor cells (mNSPCs) that again showed robust colour exchange after addition of Cre recombinase (Figure 2D-E). Probing for stability of tagged histones in non-dividing cells (Toyama et al., 2019), we used H3.1-iCOUNT mNSPCs analysed by FACS over a time course of one week post Cre, which showed a cessation of the colour exchange when cells became quiescent (by adding BMP4; Mira et al., 2010), or differentiated into neurons (by withdrawing growth factors (GF); Figure S2A-D). Finally, we designed a multiple inducible cell division counter (miCOUNT) that allows a first, Cre-dependent colour switch (red to green), followed by a second, Flipase (Flp)-dependent colour switch (green to blue; Figure 2F). mESCs expressing H3.1-miCOUNT were transduced with Cre and tamoxifen (TAM)-inducible Flp, and time-lapse imaging revealed first a switch from red (tdTomato) to green (mNeonGreen) fluorescence, followed by a colour exchange from green to blue (Cerulean) fluorescence after addition of TAM (24h after start of imaging; Figure 2G-H and Movie S2). Taken together, we show that the iCOUNT system, using different tagged proteins in different cell types, reliably reports the number of previous cell division events *in vitro*.

**Figure 2:**
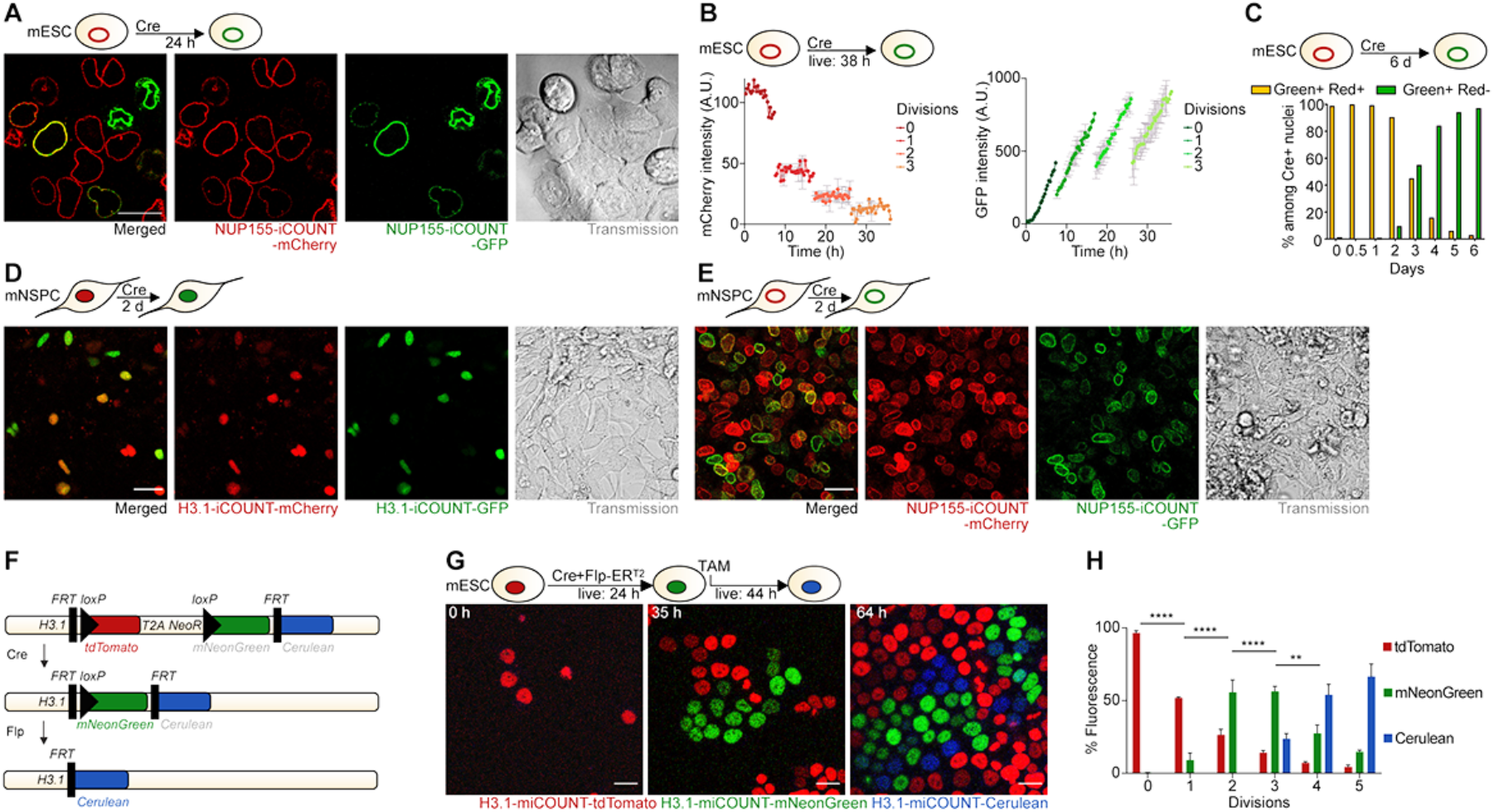
iCOUNT and miCOUNT signal in diverse mouse cell types. (A) Expression of mCherry-(red) and GFP-tagged (green) nuclear pore proteins NUP155 in mouse ESCs 24h after Cre-mediated recombination. (B) Measured changes in fluorescence intensities for red (mCherry) and green (GFP) NUP155 of single cells (mean ± SD). (C) Quantification of measured changes in fluorescence intensities for red and green NUP155-iCOUNT expressing cells using FACS. (D) Expression of mCherry-(red) and GFP-tagged (green) histones in mouse NSPCs 2d after Cre-mediated recombination. Left panel shows merged signal. (E) Expression of mCherry-(red) and GFP-tagged (green) NUP155 in mouse NSPCs 2d after Cre-mediated recombination. Left panel shows merged signal. (F) miCOUNT genetic knock-in design. (G) Selected time points of live imaging of miCOUNT-targeted cells for 68h. (H) Quantification of fluorescence intensities for red (tdTomato), green (mNeonGreen) and blue (Cerulean) histones of miCOUNT-targeted mouse ESCs grouped by cell divisions. Note the initial increase in newly synthesized green histones in miCOUNT-targeted cells after Cre-mediated recombination that is then replaced by newly synthesized blue histones after induction of Flp. \*\**p* < 0.01, \*\*\*\**p* < 0.0001. Scale bars represent 20μm.

### In vivo visualization of individual cell division histories using iCOUNT

To make use of the iCOUNT system for the analysis of cell division histories *in vivo*, we generated H3.1-iCOUNT knock-in mice. First, the iCOUNT mouse was crossed to a mouse line ubiquitously expressing inducible Cre recombinase (ROSA26:Cre-ER^T2^). Mouse embryos were analysed at embryonic day (E) 11.5 or E15 after Cre was induced using TAM injection at E9.5 or E13.5, respectively (Figure 3A and Figure S3A). Green cells were observed in all developing organs including the developing brain, liver, gut, somites, eye and skin (Figure 3A-B). Fluorescently labelled histones were symmetrically segregated during mitosis and green/red ratios were preserved after fixation and antibody-based amplification (Figure S3B-C). Focusing on the developing neocortex at E11.5, columns of green cells were observed, and within those the percentage of green fluorescence (green/total fluorescence) was increased in actively dividing cells (KI67+; Figure 3C-D), indicating that iCOUNT signal reports cell division events *in vivo*.

**Figure 3:**
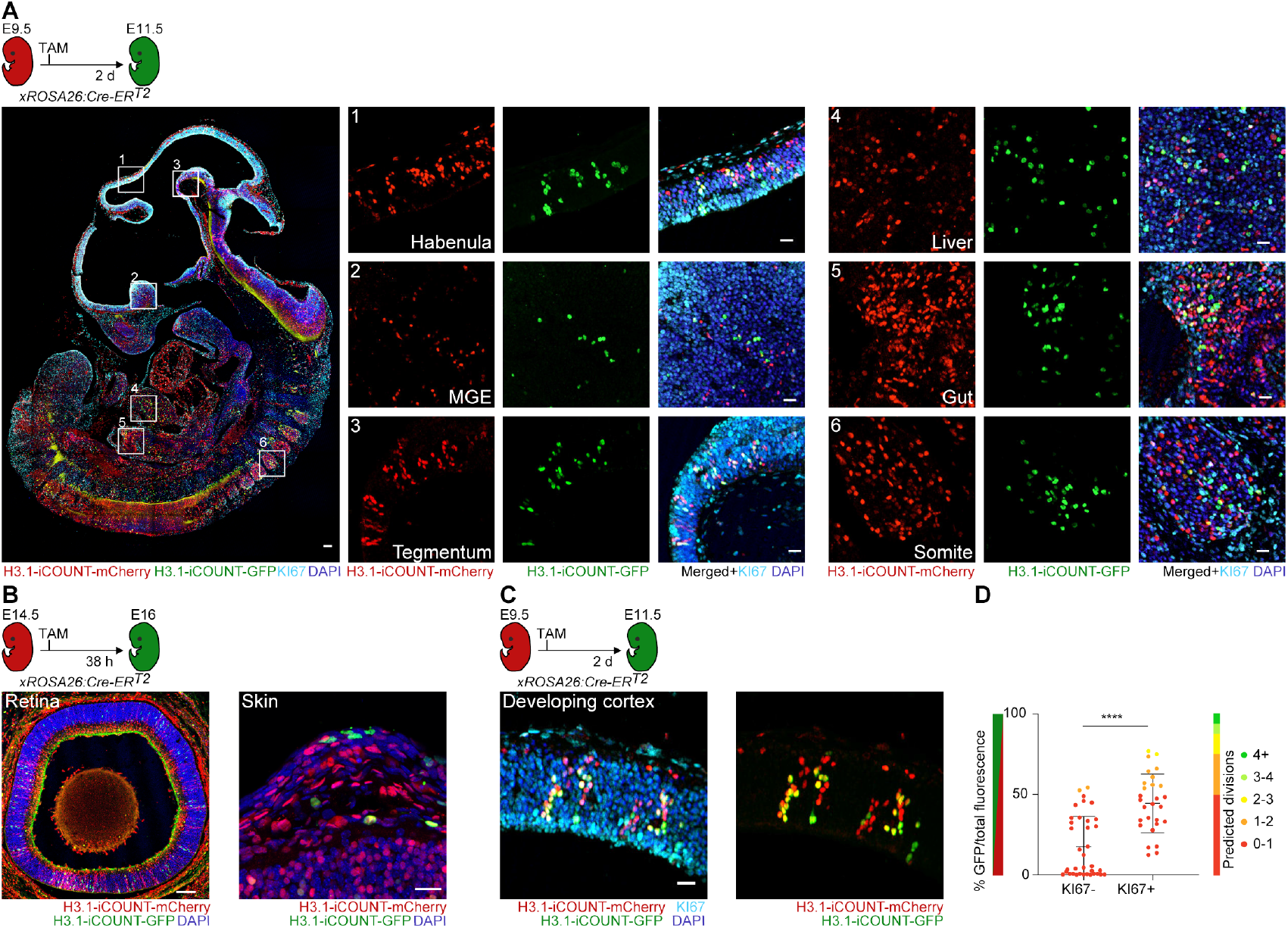
iCOUNT visualizes cell division history in embryonic tissues. Overview of iCOUNT mouse at E11.5 (recombination using TAM at E9.5). Right panels show mCherry (red), GFP (green), KI67 (light blue) signal in tissues indicated. Overview of the developing retina and skin in iCOUNT embryos induced with TAM at E14.5 and analysed at E16. (C) iCOUNT-targeted cells in the developing cortex. (D) Graph shows quantification of green/total fluorescence and predicted division numbers of cells grouped by KI67 expression. Nuclei were counterstained with DAPI (blue). \*\*\*\**p* < 0.0001. Scale bars represent 100μm in A, B (retina), and 20μm in A (right panels), B (skin) and C.

Next, Cre recombinase was induced at E14.5 and the ratio of red/green fluorescence was analysed in cells within different areas of the developing cortex and at different time points after Cre induction (E15.5, E16 and E16.5; Figure 4A). Strikingly, the percentage of green fluorescence increased with time after Cre induction with an initial increase in the ventricular and subventricular zone (VZ/SVZ), where radial glia cells are constantly dividing, and later in the intermediate zone (IZ), and last in the cortical plate (CP; Figure 4B and Figure S4A-B) (Gotz and Huttner, 2005). These data suggest that radial glia cells divide between zero and >4 times within 50h, generating neurons with gradually increasing fractions of green fluorescence that migrate into the IZ and later to the CP. These findings are consistent with previous estimates of cell cycle and migration durations (Figure 4C and *Theory* in Methods) (Gao et al., 2014). We next used the iCOUNT system to test the hypothesis that the same radial glia cells first generate deep layer and later upper layer neurons within the cortex (Lui et al., 2011). Cre was induced at the start of neurogenesis (E11.5) and cortices were analysed after completion of cortical neurogenesis (E19.5). iCOUNT data revealed an increase in the percentage of green fluorescence in cells within the upper layers compared to deep layers (Figure 4D-E). However, we observed a substantial overlap between the two layers (Figure 4F). iCOUNT identifies overall similar numbers compared to previously reported mathematical modelling data derived from MADM-based clonal data (Figure 4G and Supplemental Computational Modeling). However, the overlap of previous cell divisions in deep and upper layer and subtle differences compared to MADM-based modelling data suggest that not all radial glia cells follow a temporally fixed program of set cell divisions, thus challenging previous snapshot-based estimates of cell division biographies of radial glia cells and their progeny in the embryonic brain (Gao et al., 2014; Gotz and Huttner, 2005).

**Figure 4:**
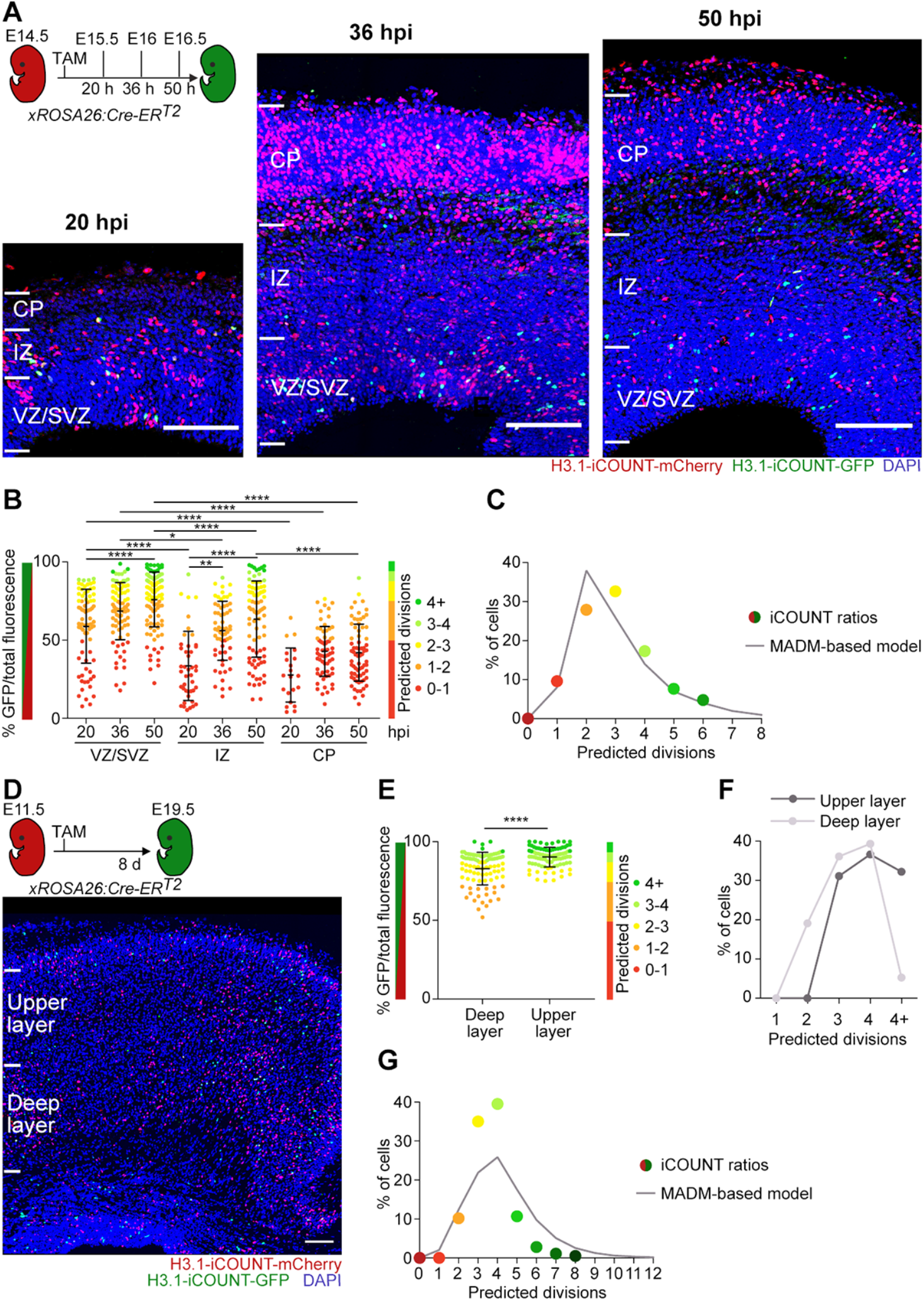
iCOUNT allows counting of previous cell divisions in the developing mouse brain. (A) iCOUNT-targeted cells in the developing cortex recombined at E14.5 and analysed at the indicated time points. (B) Quantification of green/total fluorescence and predicted division numbers of cells in the VZ/SVZ, IZ, CP at different time points. (C) Percentage of cells with iCOUNT-predicted cell divisions compared to MADM-based modelling data (Gao et al., 2014) in the VZ/SVZ 50h post induction. (D) iCOUNT-targeted cells in the developing cortex recombined at E11.5 and analysed at E19.5. (E) Quantification of green/total fluorescence and predicted division numbers of cells in the deep and upper cortical layers. (F) Percentage of cells with predicted cell division numbers in upper (dark grey) and deep (light grey) cortical layer. (G) Percentage of cells with iCOUNT-predicted cell divisions compared to MADM-based modelling data (Gao et al., 2014) in cortical layers (E11.5 to E19.5 chase). Nuclei were counterstained with DAPI (blue). \**p* < 0.05, \*\**p* < 0.01, \*\*\*\**p* < 0.0001. Scale bars represent 100μm.

To use the iCOUNT in adult mice, we examined the expression of H3.1-iCOUNT in different organs and found mCherry positive cells in all tissues analysed (shown are brain, gut, liver, kidney; Figure 5A). Crossing H3.1-iCOUNT mice to ROSA26:Cre-ER^T2^ mice allowed the analysis of red/green cells in different organs. Using FACS analysis two days after Cre induction we observed red/green cells for example in the brain, bone marrow and skin (Figure 5B), indicating the broad usability of H3.1-iCOUNT mice. To focus on adult NSPCs, iCOUNT mice were crossed with mice expressing inducible Cre recombinase under the Gli1 promoter (Gli1:Cre-ER^T2^) leading to expression of inducible Cre selectively in adult NSPCs (Ahn and Joyner, 2005). Two weeks after TAM-induced Cre-mediated recombination, red/green cells were mostly observed in two regions, the dentate gyrus (DG) and the subventricular zone, with only few cells labelled in the hypothalamus, in agreement with previous literature reporting neurogenic zones in the adult brain (Figure 5C) (Braun and Jessberger, 2014). Using iterative immunostaining technology (4i), we performed immunofluorescent labelling of different marker proteins besides the iCOUNT colours in red and green to phenotype recombined cells (Figure 5D and Figure S5A-B (Gut et al., 2018). We used antibodies detecting HOPX, SOX2, GFAP, DCX, IBA1, NEUN, KI67, S100β and OLIG2 on whole brain sections to distinguish different cell types in the DG: radial glia-like (R) cells (HOPX+, SOX2+, GFAP+, with radial process); non-radial glia-like (NR) cells (HOPX+, SOX2+, without radial process), dividing NR cells (HOPX+, SOX2+, KI67+), newborn neurons (DCX+), mature neurons (NeuN), astrocytes (SOX2+, GFAP+, S100β+), oligodendrocytes (OLIG2+) and microglia (IBA1+) (Figure S5A-B) (Berg et al., 2019). As expected, most iCOUNT green cells were categorized as R, NR cells or newborn neurons (Figure 5D). Within those, R cells showed the lowest percentage of green fluorescence that increased in NR cells and that was further elevated in actively dividing (KI67+) NR cells and newborn neurons (Figure 5E). We used iCOUNT to predict cell division histories of individual cells derived from R cells in the adult neurogenic lineage (Figure 5F). Strikingly, iCOUNT-derived numbers closely matched cell division estimates of R, NR cells and neurons using chronic *in vivo* imaging approaches two weeks after Gli1-dependent reporter expression (Figure 5F), thus cross-validating adult brain intravital imaging and iCOUNT-based technologies (Pilz et al., 2018). Together, the iCOUNT data confirm that R cells divide few times in the adult brain, whereas NR cells substantially amplify the pool of neural progenitor cells that subsequently differentiate into neurons originating from a range of mother cell divisions (Bonaguidi et al., 2011; Pilz et al., 2018).

**Figure 5:**
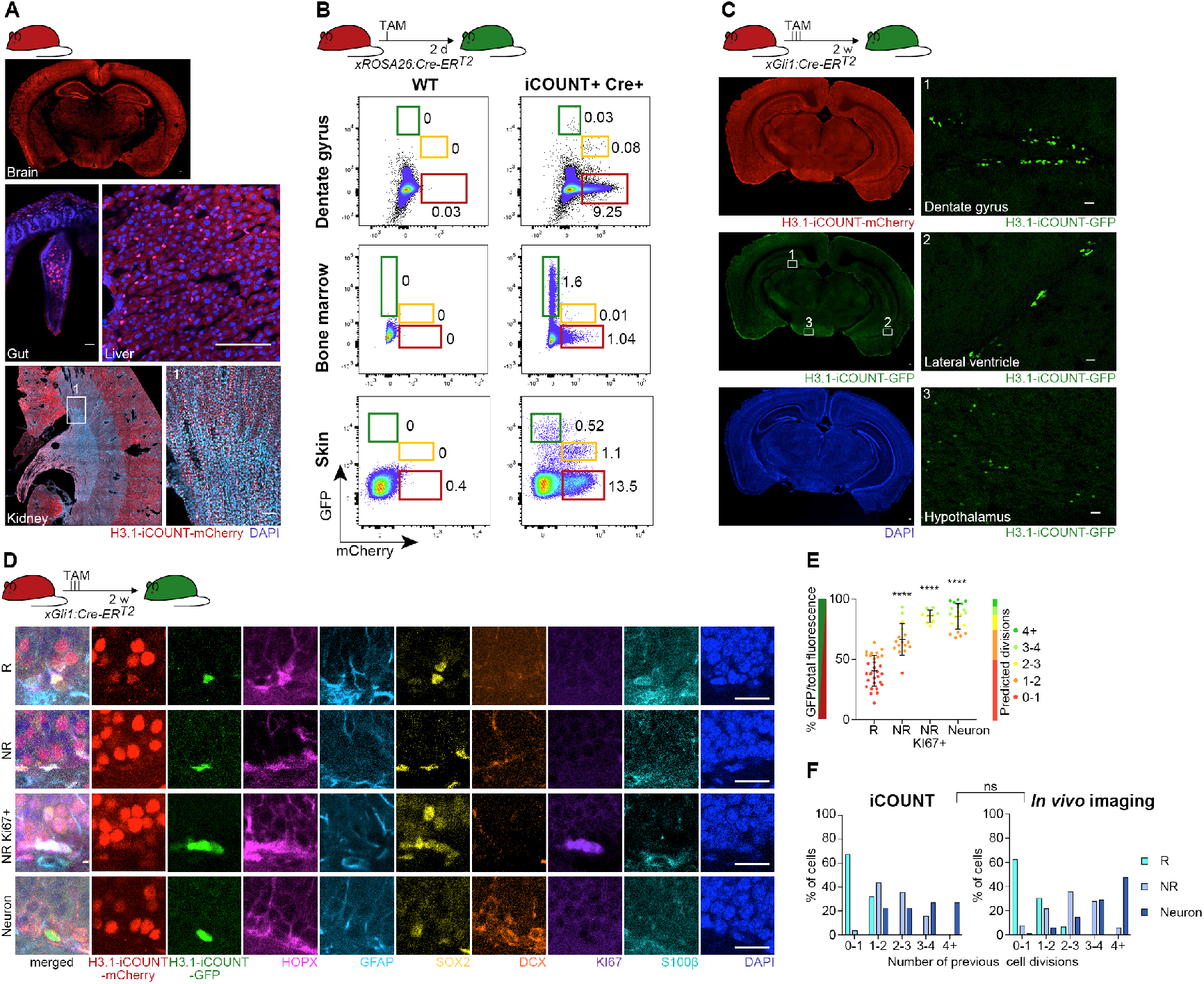
iCOUNT signal in adult mouse tissues. (A) Overview of an adult iCOUNT mouse with expression of the tagged H3.1 in diverse tissues as indicated. (B) FACS analyses of cells with distinct red/green histone ratios in diverse tissues as indicated. Left panels show controls, right panels show cells expressing newly synthesized (green) histones after inducible Cre-based recombination. (C) Conditional recombination in adult NSPCs identifies iCOUNT-targeted cells in the adult brain. Left panels show overview, boxed areas are shown in higher magnification (right panels). (D) 4i-based phenotyping of Cre-targeted cells in the adult hippocampus using a panel of protein markers as indicated reveals radial glia-like NSPCs (R), non-radial glia-like NSPCs (NR), actively dividing NR (KI67+) and neurons. (E) Quantification of green/total fluorescence and predicted division numbers of cells in the adult hippocampus 2 weeks after Cre-based recombination. (F) Number of previous cell divisions based on iCOUNT and chronic intravital imaging. Note the comparable distribution among R, NR and neurons. Nuclei were counterstained with DAPI (blue). \*\*\*\**p* < 0.0001, ns, not significant. Scale bars represent 100μm in A, C and 20μm in C (right panels), D.

To study individual cell division histories in human cells, we first tagged H3.1 with miCOUNT in human embryonic stem cells (hESCs) and followed single cells using time-lapse microscopy after Cre, observing the predicted colour exchange also in human cells (Figure 6A-B). Thus, we generated human forebrain-regionalized organoids from miCOUNT hESCs (Qian et al., 2016). Organoids were first treated with Cre for 24h and then live whole organoid imaging was performed for 54h. Visualization of single cells within an intact organoid revealed the expected colour exchange and symmetric segregation of miCOUNT tagged histones (Figure 6C, Figure S6A-B, and Movie S3). Next, we performed 4i on organoids and found the highest percentage of green fluorescence in the inner part of cortical units compared to the middle, outer parts and newly generated neurons (Figure 6D-E and Figure S6C). To test for dual colour switch in hESCs, ER^T2^-Cre-ER^T2^ expressing NUP155-miCOUNT hESCs were generated, treated for two days with TAM and then electroporated with Flp recombinase. A colour exchange from red (tdTomato) to green (GFP) and then from green to blue (tagBFP) was observed both by fluorescent imaging and FACS (Figure S6D-E).

**Figure 6:**
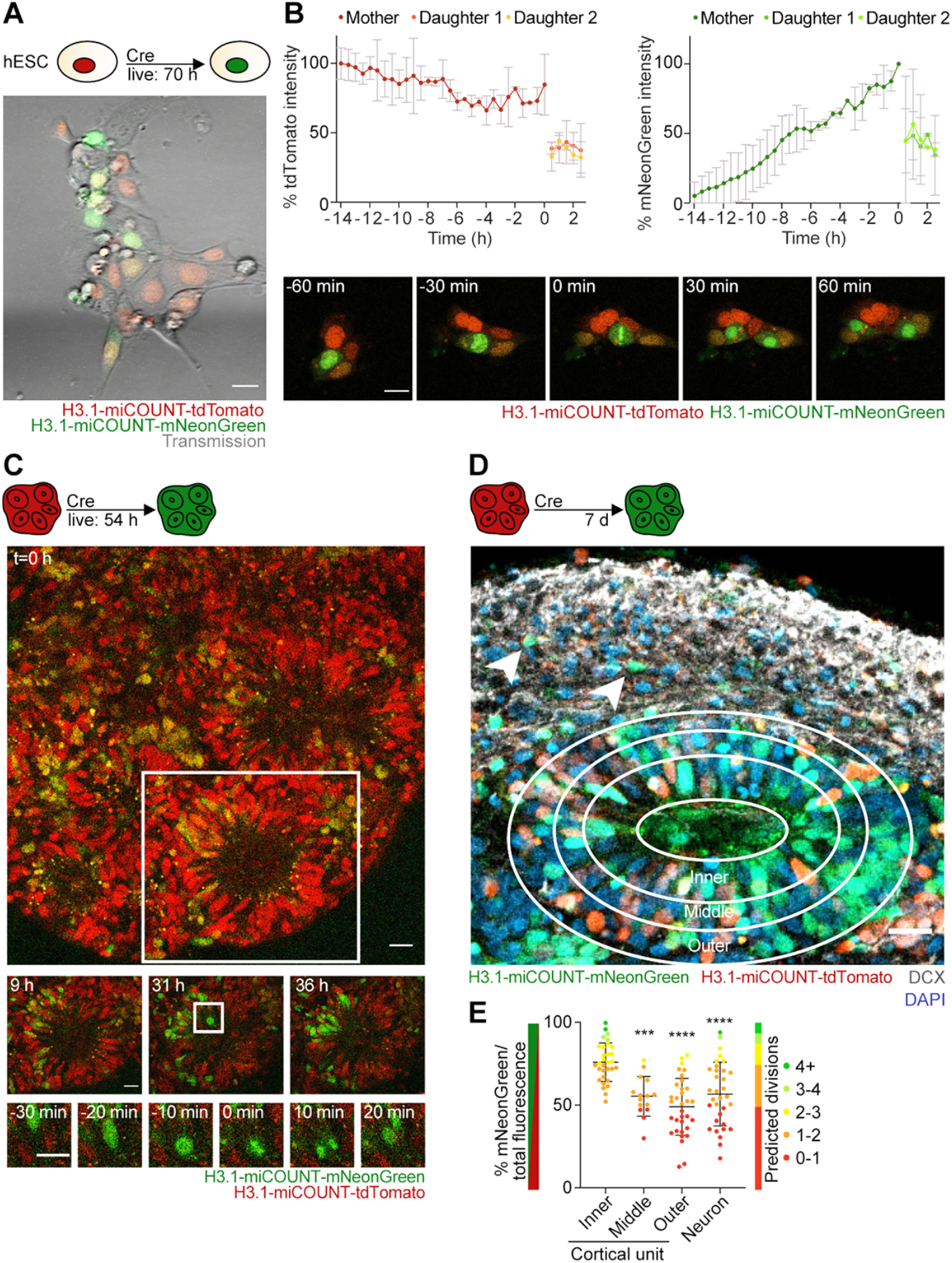
miCOUNT signal in human forebrain organoids. (A) Selected time point of live imaging for 70h after induction with Cre, showing tdTomato (red) and mNeonGreen (green)-tagged histones in human ESCs. (B) Graph shows measured changes in fluorescence intensities for red (tdTomato) and green (mNeonGreen) histones of single cells aligned by the time of division (mean ± SD). Lower panel shows images of a dividing human ESC with high temporal resolution. (C) Selected time points of live imaging of human miCOUNT expressing cells in a forebrain organoid for 54h. Middle panels show high magnifications of the cortical unit boxed in the top panel. Lower panels show high magnifications of a dividing progenitor boxed in the middle panel. (D) Micrograph of miCOUNT cells 7 days after Cre-based recombination shows increase in mNeonGreen (green) histones in the inner parts of the cortical unit (circles depict analysed areas). Neurons are stained with DCX (white, arrow heads point towards examples of neurons). (E) Quantifications of green/total fluorescence and predicted division numbers of cells in human organoids in distinct areas of a cortical unit as indicated. Nuclei were counterstained with DAPI (blue). \*\*\**p* < 0.001, \*\*\*\**p* < 0.0001. Scale bars represent 20μm.

### Molecular consequences of previous cell division events in mouse and human NSPCs

We next aimed to identify the molecular consequences of previous cell divisions on individual cells using single cell RNA-sequencing (scRNA-seq). First, we used FACS to sort single cells with different fractions of red/green fluorescence (Figure S7A) of human miCOUNT organoids four and seven days after Cre injections, followed by Smart-Seq2-based scRNA-seq (Picelli et al., 2014). In line with previous organoid scRNA-seq experiments, t-distributed stochastic neighbour embedding (t-SNE) revealed four clusters, which were classified as dividing and non-dividing NSPCs, and immature and mature neurons, based on differential expression of known marker genes (Figure 7A) (Camp et al., 2015). Within each cluster, cells of different iCOUNT red/green ratios were present, allowing us to compare their gene expression profiles (Figure 7B). In parallel, cells from mouse developing cortices were collected at E15, 38h post-induction of Cre, and FACS sorted based on their iCOUNT ratio (Figure S7B). Again, t-SNE analysis revealed four major clusters: apical and basal progenitors, as well as immature and mature neurons, based on expression of marker genes (Figure 7C), with cells containing different red/green ratios in all clusters (Figure 7D).

**Figure 7:**
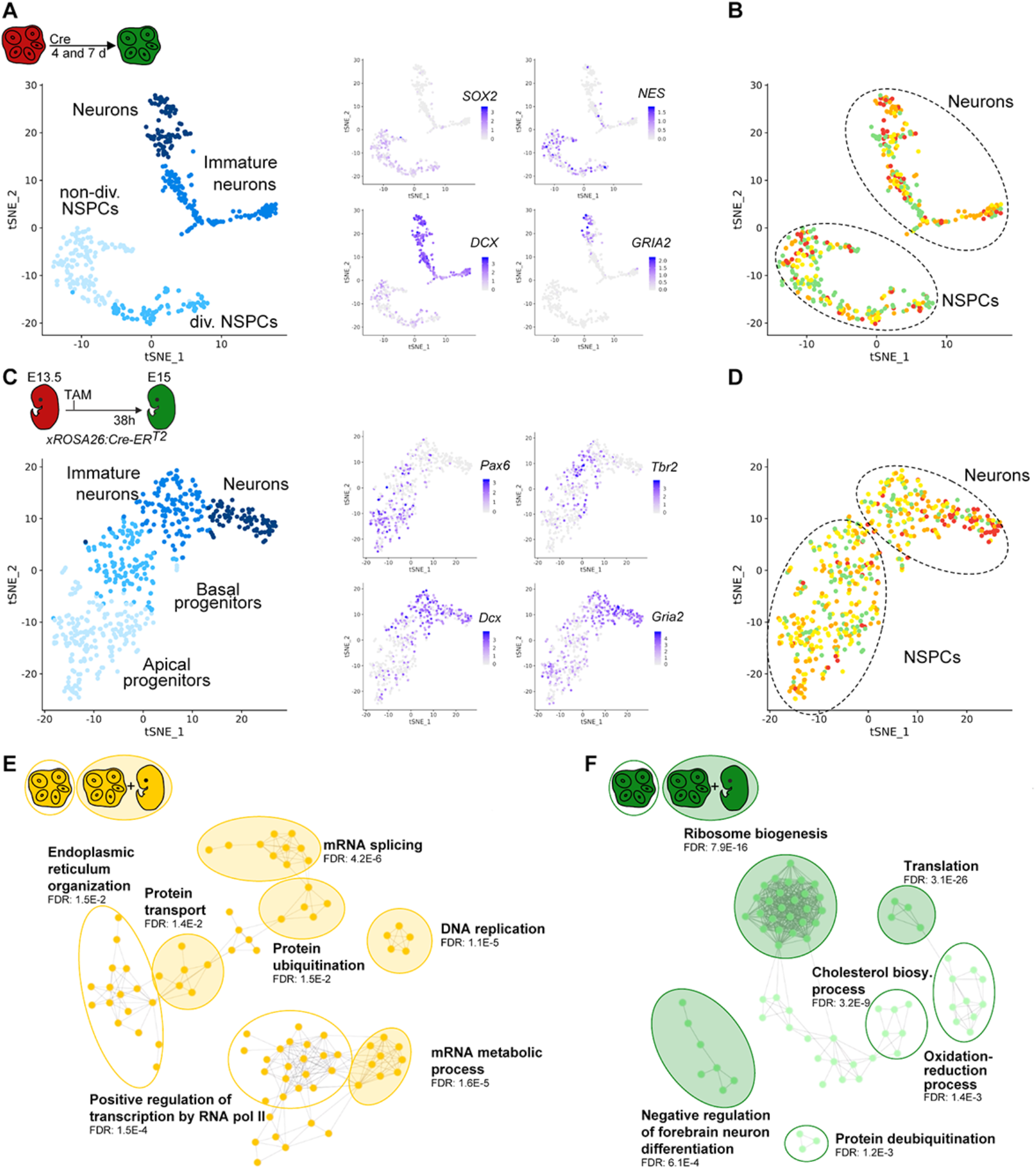
Molecular consequences of previous cell divisions in mouse and human NSPCs. (A) scRNA-seq of human forebrain organoids identifies expected clusters of cells (left panel shows tSNE) with representative marker genes imposed on identified cluster (right panels). (B) Clustering of iCOUNT-targeted human cells into NSPCs and neurons. Cells with different green/total fluorescence ratios are indicated on the tSNE plot. (C) scRNA-seq of developing mouse cortex identifies expected clusters of cells (left panel shows tSNE) with representative marker genes imposed on identified cluster (right panels). (D) Clustering of iCOUNT-targeted mouse embryonic cells into NSPCs and neurons. Cells with different green/total fluorescence ratios are indicated on the tSNE plot. (E) GO enrichment analyses of cells with low (orange) cell division history within the NSPC cluster of human organoids. GO terms shown in shaded circles were shared between human organoids and developing mouse cortex. (F) GO enrichment analyses of cells with high (green) cell division history within the NSPC cluster of human organoids. GO terms shown in shaded circles were shared between human organoids and developing mouse cortex.

Gene expression profiles of cells within the NSPC cluster of human organoids (Figure 7B) were used to compare orange (low dividers) and green cells (high dividers). We identified 241 genes upregulated in orange and 120 genes in green cells (Table S1-2). Gene ontology (GO) enrichment analysis revealed different clusters, of which many were also found in the GO enrichment analysis of the developing mouse cortex scRNA-seq data (filled circles; Figure 7E-F and Table S3). Genes enriched in orange iCOUNT cells within the NSPC clusters were, for example, involved in DNA replication (Figure 7E), whereas genes enriched in green cells were involved in ribosome biogenesis (Figure 7F). Thus, we here discovered that cells with different individual cell division histories are distinct in their gene expression profiles.

## Discussion

Recently, substantial progress has been made to derive full lineage tree information from individual cells using barcoding techniques (Baron and van Oudenaarden, 2019; Fuentealba et al., 2015; Gao et al., 2014; Kalhor et al., 2018; Mayer et al., 2015; McKenna et al., 2016). However, potential intermediate steps are inherently hidden in these approaches, making it challenging to infer information such as which cells divided how many times, and how many cells were born but later lost due to cell death. Several approaches have tried to overcome these fundamental problems. One strategy is to load cells with a dye that becomes diluted with consecutive cell divisions, such as carboxy-fluorescein-succinimidyl-ester (CFSE) (Lyons and Parish, 1994; Takizawa et al., 2011). This approach allowed for the generation of fundamentally important insights, for example in the understanding of hematopoietic stem cell (HSC) behaviour (Takizawa et al., 2011; Vannini et al., 2016). However, CFSE loading requires the isolation and subsequent re-transplantation of somatic stem cells, which will substantially affect the cellular properties of analysed cells and also renders its application only possible to few selected somatic stem cell niches, such as HSCs. Other approaches used in mice are based on transgenesis-mediated loading of fluorescently labelled histones, such as histone H2B fused to GFP (Bernitz et al., 2016; Foudi et al., 2009; Qiu et al., 2014; Tomasetti and Vogelstein, 2015; Tumbar et al., 2004). However, it has been shown that H2B-GFP is exchanged even without cell division events, alters cellular properties due to artificial histone overexpression, and affects nucleosome structure and function, preventing its compatibility with transcriptomics. (Challen and Goodell, 2008; Meeks-Wagner and Hartwell, 1986; Ricci et al., 2015; Toyama et al., 2019; Toyama et al., 2013; Tumbar et al., 2004). Furthermore, the H2B-GFP strategy turned out to be partially leaky and thus not useful in a variety of systems and does not allow for calibration of cell division events for each individual cell as only one fluorescent signal (i.e., its dilution) is analysed (Challen and Goodell, 2008; Tumbar et al., 2004). In contrast, iCOUNT is inserted into the endogenous locus and does not lead to overexpression of the gene, exemplified here by using H3.1 and NUP155. Further, iCOUNT-based strategies allow for the analysis of fluorescent ratios, thus eliminating variations based on differential expression, cell size, and imaging conditions. Importantly, iCOUNT can be combined with clonal lineage tracing approaches, in contrast to previously described H2B-GFP-based approaches. However, H3.1 tagging may cause mosaic expression of the RITE cassette, as there are 9 different genes encoding for the histone-variants H3.1 and H3.2 within the *HIST1* gene cluster on chromosome 13 (Wang et al., 1996). Indeed, we observed mosaic expression and mice that had the RITE cassette correctly inserted (verified via genomic PCR) but did not express the transgene.

We show the accuracy of iCOUNT using *in vitro* time-lapse imaging and by comparing cell division estimates obtained from adult NSPCs using iCOUNT to data that were obtained by chronic intravital imaging. The iCOUNT data matched well with the MADM-based modelling. We thereby demonstrate the usability of the iCOUNT system both *in vitro* and *in vivo*. Comparing the division history of neurons in the deep and upper layer of the developing mouse cortex revealed that indeed, radial glia cells generating deep layer neurons divided less times compared to those generating upper layer neurons, supporting the current model (Gao et al., 2014). However, we observed a substantial overlap between deep and upper layer neurons.

These findings indicate that not all radial glia cells first produce deep and later upper layer neurons but rather suggests the existence of different radial glia cells being active at different times during development. Further characterization using iCOUNT embryos at different time points will render important new insights into the division pattern of single radial glia cells during cortex development in mice.

Important insights into the division potential and distinct responsiveness of stem cells have been derived from intravital imaging (Barbosa et al., 2015; Gurevich et al., 2016; Lo Celso et al., 2009; Pilz et al., 1998; Ritsma et al., 2014; Rompolas et al., 2012; Xie et al., 2009). However, chronic imaging is not possible in a variety of stem cell niches and not easily compatible with methods to isolate cells for molecular phenotyping. In contrast, we were able to isolate hundreds of cells with individual division histories from various organs simultaneously, and analyse their molecular profiles using scRNA-seq. As a proof-of-principle we performed scRNA-seq of mouse developing cortex and human forebrain organoids. We discovered networks of differentially expressed genes depending solely on the cell division history of the cells. Importantly, several of them were conserved between mouse and human cells. Future research will manipulate those networks to probe their relevance for cell division potential of stem cells. Furthermore, comparing the expression profiles of different iCOUNT-coloured cells derived from other somatic stem cell compartments will render both stem cell specific and global effects of an individual cell’s division history. This will greatly broaden our understanding of behavioural and molecular consequences of previous cell divisions of different cell types in mammalian stem cell niches.

The miCOUNT approach will not only allow the exact counting of more divisions but also allows for the visualization of stem cell division patterns during two different time points, for example during development and in the adult organism. Together, determining the cell division biography of individual cells throughout life span and its molecular consequences will further our understanding of stem cell behaviour in health and disease.

## Acknowledgements

We thank Y. Barral for conceptual input, C. Rinaldo, C. Paulson, C. Trindade Antunes, V. Korobeynyk, J.D. Cole for experimental help, S. Bottes and G. A. Pilz for sharing data, E. Yángüez López-Cano from the Functional Genomics Center Zurich (UZH /ETHZ) and the Cytometry Facility of UZH. A.D.L. was supported by a Forschungskredit of UZH and the Novartis foundation. The authors declare no competing financial interests. This study was supported by the European Research Council (STEMBAR to S.J.), the Swiss National Science Foundation (BSCGI0_157859 to S.J.), and the Zurich Neuroscience Center (ZNZ).

## Author contributions

A.D.L. developed the concept, performed experiments, analysed data, and wrote the manuscript. B.N.J. performed experiments, analysed data, and revised the manuscript. T.L, S.E.C., L.N.R., M.K. performed experiments, analysed data. B.D.S. analysed data and revised the manuscript. S.J. developed the concept and wrote the manuscript.

## STAR★Methods

### Key Resources Table

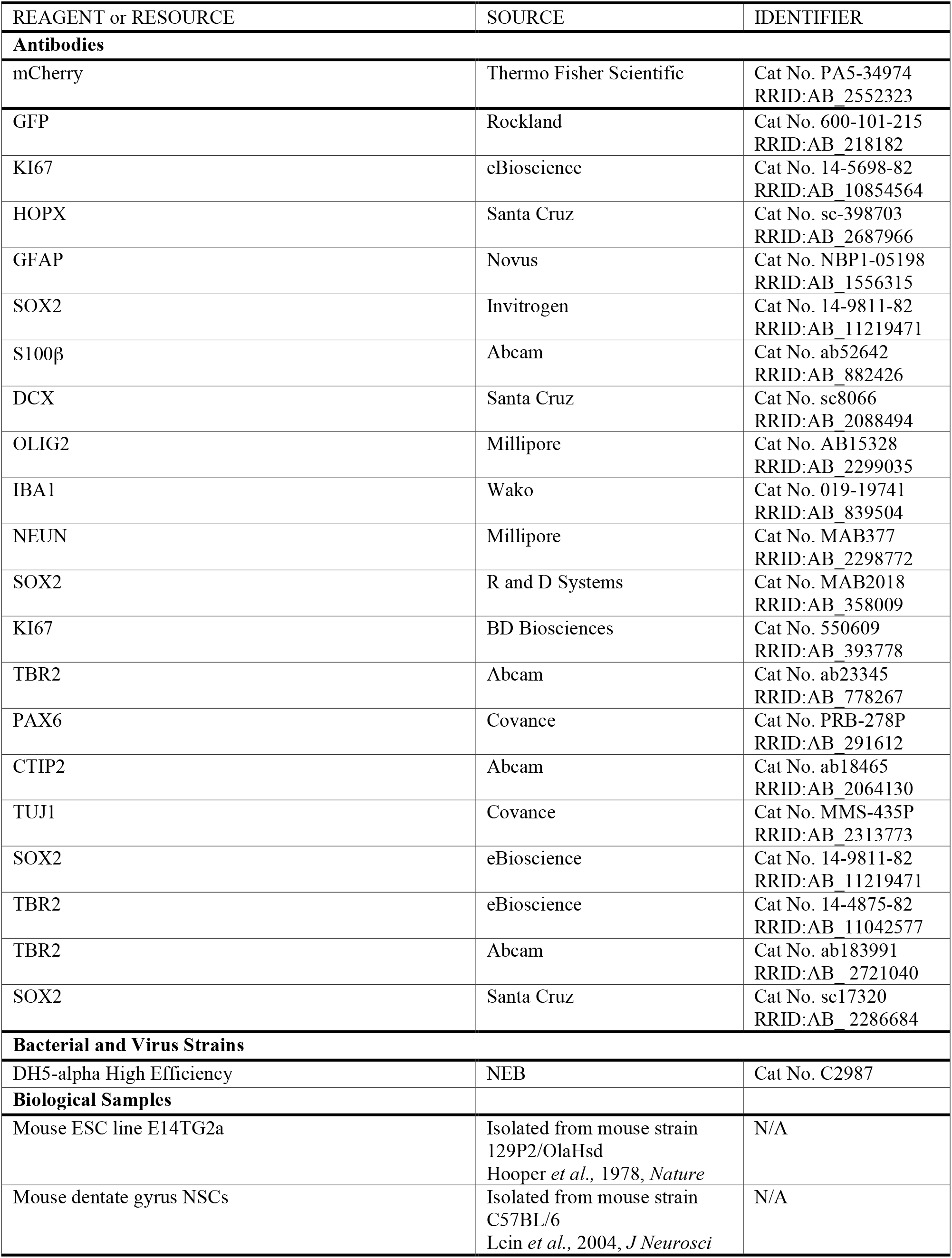
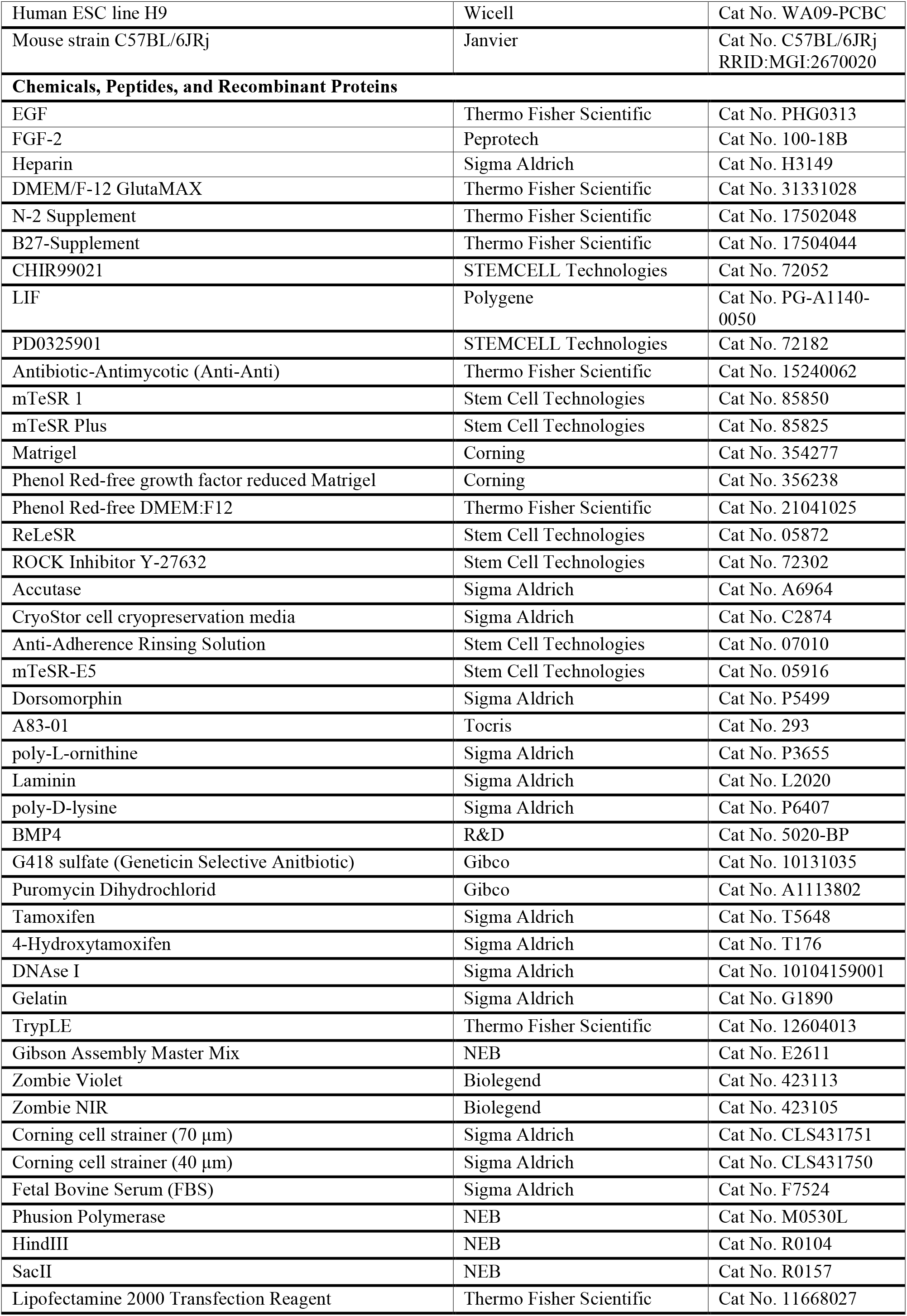
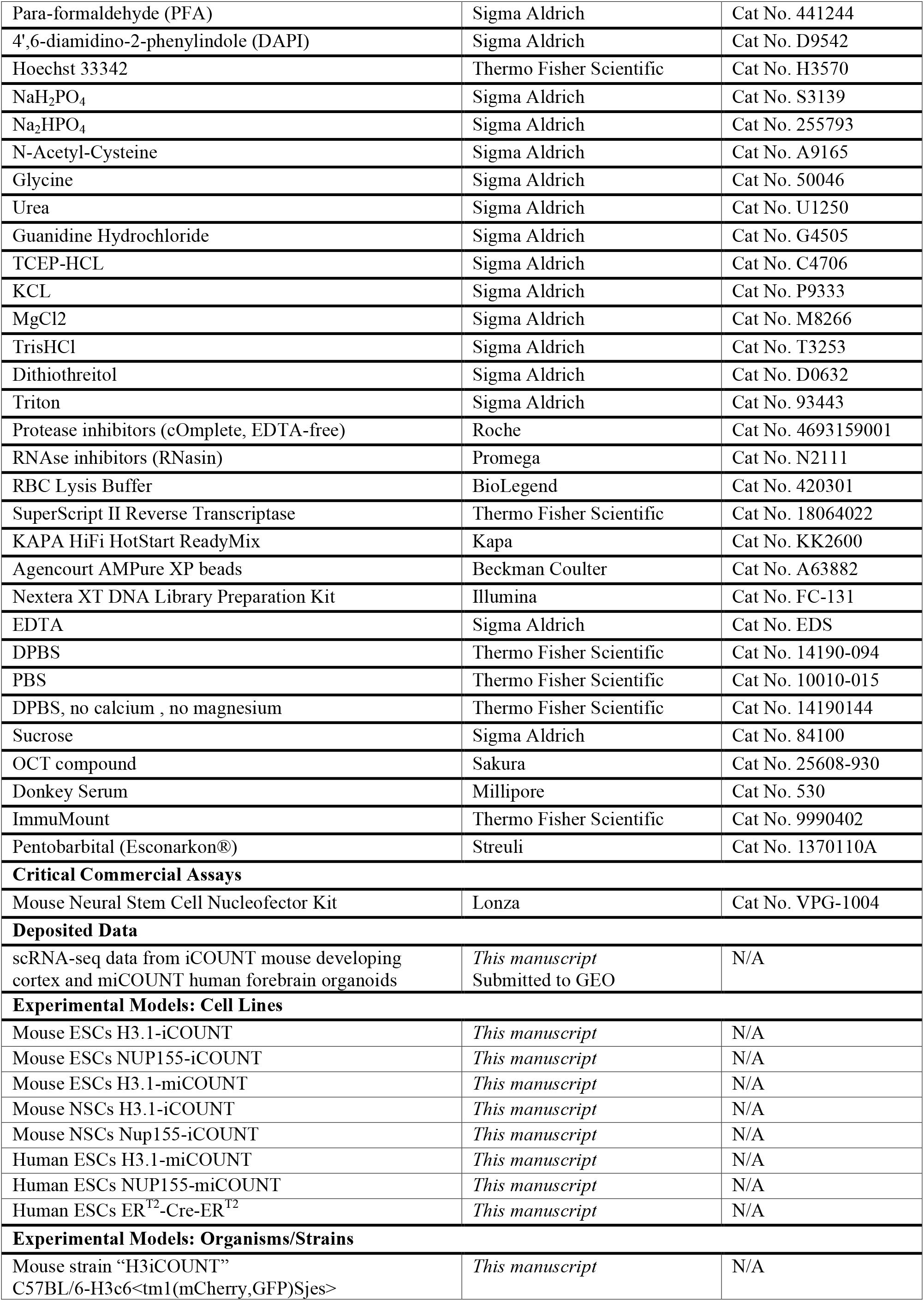
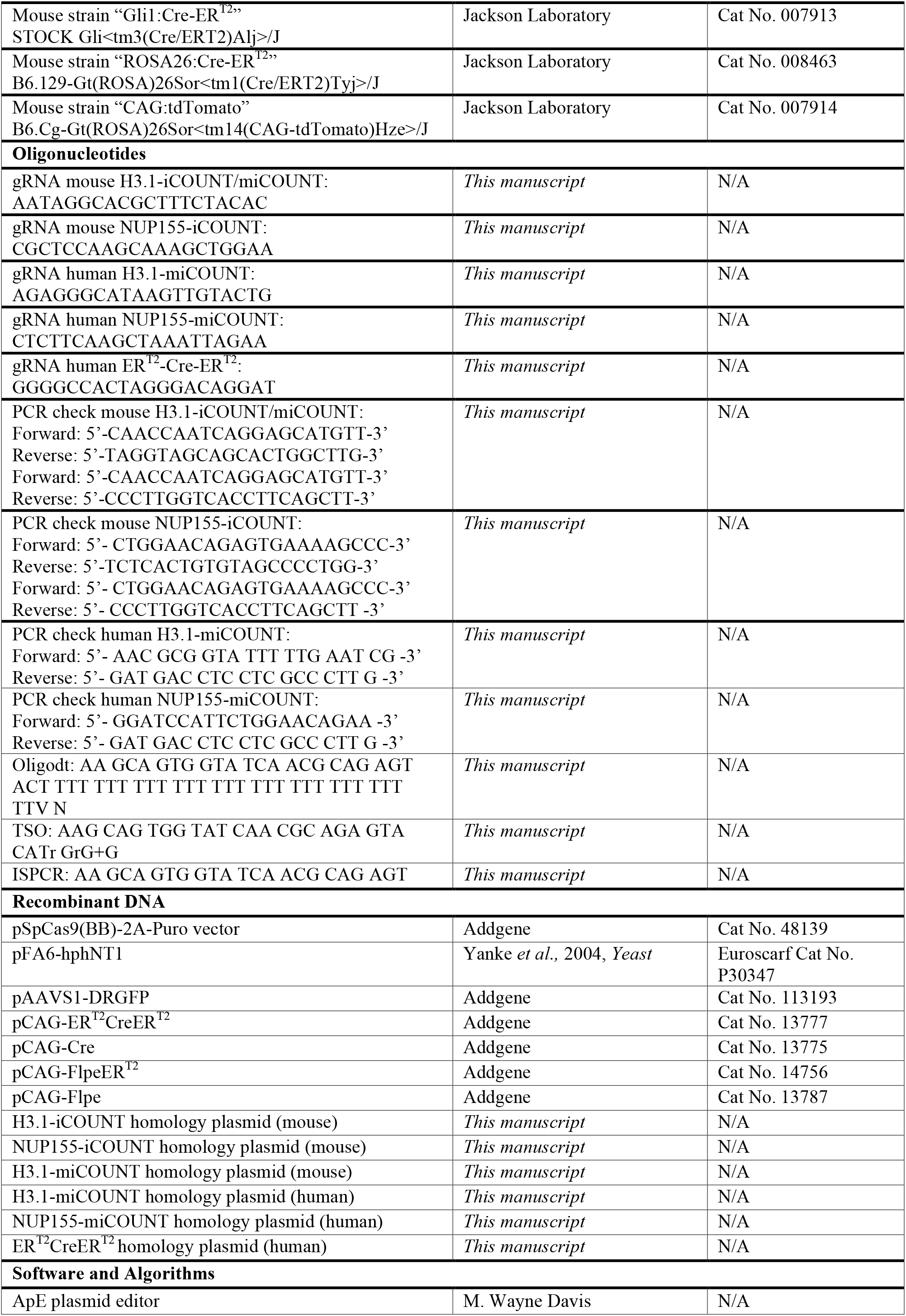
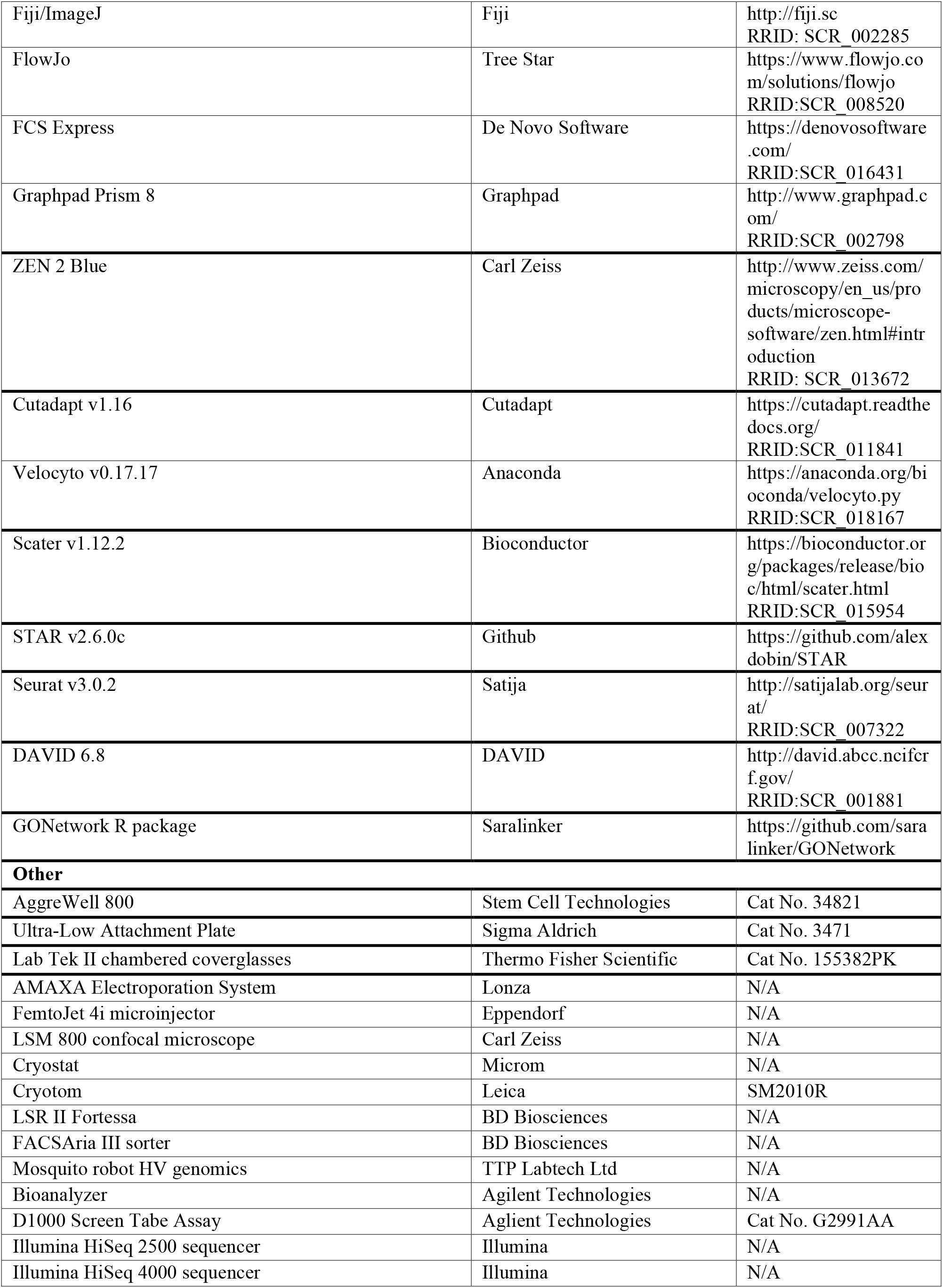

## Lead contact and materials availability

Further information and requests for resources and reagents should be directed to and will be fulfilled by the Lead Contact, Sebastian Jessberger. iCOUNT/miCOUNT mouse and human ESCs and iCOUNT-mice will be provided directly. Materials will be provided upon execution of a suitable Materials Transfer Agreement.

## Experimental model and subject details

### Stem cell cultures

Mouse embryonic stem cells (mESCs; E14TG2a; Hooper et al., 1987) were cultured in DMEM/F12 GlutaMAX medium (Thermo Fisher Scientific) supplemented with N2 and B27 (Thermo Fisher Scientific), antibiotics (Anti-Anti, Thermo Fisher Scientific), LIF (1000U/ml, Polygene), and 2i (Silva et al., 2008), PD0325901 (1μM, STEMCELL Technologies) and CHIR99021 (3μM, STEMCELL Technologies). mESCs were cultured in flasks coated with 0.2% gelatine (Sigma-Aldrich). Plasmids were introduced into cells using lipofection (Lipofectamine 2000, Thermo Fisher Scientific).

Neural stem/progenitor cells (NSPCs) derived from the hippocampus of adult C57BL/6 animals were cultured in DMEM/F12 GlutaMAX medium supplemented with N2, B27, antibiotics, EGF (20ng/ml, Thermo Fisher Scientific), FGF-2 (20ng/ml, PeproTech) and Heparin (5mg/ml, Sigma-Aldrich). Plasmids were introduced using the AMAXA electroporation system (Lonza) as described in (Wegleiter et al., 2019). NSPCs were differentiated into neurons by growth factor withdrawal (no addition of EGF and FGF-2) and quiescence was induced by exchanging EGF with BMP4 (Knobloch et al., 2017; Mira et al., 2010) (50ng/ml, R&D Systems).

Human embryonic stem cells (hESCs; H9 (Thomson et al., 1998), Wicell) were grown in feeder-free conditions in the absence of antibiotics. H9 hESCs were maintained in mTeSR 1 or mTeSR Plus (Stem Cell Technologies) on hESC qualified Matrigel (Corning) coated plates. Routine passaging was performed with ReLeSR (Stem Cell Technologies) to promote stemness; post-passaging, cells were maintained for 24h in media containing Y-27632 (10μM, Stem Cell Technologies). To produce single cell suspensions, hESCs were passaged with Accutase (Sigma-Aldrich). For long-term storage, hESCs were kept below −170°C in CyroStore CS10 (Sigma-Aldrich). To introduce plasmid DNA, hESCs were pretreated with mTeSR 1 or Plus containing Y-27632 for at least 2h to improve cell survival. Two million cells were resuspended in Nucleofector V (Lonza) and electroporated with an AMAXA electroporation system using program A-23 according to the manufacturer’s guidelines. All cells were kept at 37°C and 5% CO_2_. All experiments using hESCs were approved by the Kantonale Ethik-Kommission (KEK) of the Canton of Zurich, Switzerland.

### Organoid culturing

Forebrain specific cerebral organoids were produced using the previously published protocol (Qian et al., 2016) with some alterations. At day 0, hESCs were passaged with Accutase and cells were resuspended in mTeSR 1 or Plus containing Y-27632. A cytometer was used to estimate the number of cells in the single cell suspension. The single cell suspension was added to pretreated wells AggreWell 800 (Stem Cell Technologies) that would result in 5000 cells per microwell. Pretreatment was performed with Anti-Adherence Rinsing Solution (Stem Cell Technologies) and handling of the AggreWell 800 plates was carried out following the manufacturer’s recommendations. At day 1, embryoid bodies were harvested from the AggreWells per the manufacturer’s protocol and transferred to a well of a 6 well Ultra-Low Attachment Plate (Sigma-Aldrich) containing mTeSR-E5 (Stem Cell Technologies) with Dorsomorphin (2μM, Sigma-Aldrich) and A83-01 (2μM, Tocris). Embryoid bodies were maintained in this medium until day 4, after which the protocol described (Qian et al., 2016) was followed. Organoids were fixed for 30min in Paraformaldehyde (PFA, 4%, Sigma-Aldrich) and were stored in sucrose (30%, Sigma-Aldrich) until further use.

### Gene tagging

Gene tagging using CRISPR/Cas9 was essentially performed as described (Ran et al., 2013). gRNAs were designed using crispr.mit.edu/ and were cloned into pSpCas9(BB)-2A-Puro (Addgene No 48139) according to the protocol (Ran et al., 2013). The following gRNAs were used:

Mouse H3.1-iCOUNT/miCOUNT: AATAGGCACGCTTTCTACAC
Mouse NUP155-iCOUNT: CGCTCCAAGCAAAGCTGGAA
Human H3.1-miCOUNT: AGAGGGCATAAGTTGTACTG
Human NUP155-miCOUNT: CTCTTCAAGCTAAATTAGAA
Human ER^T2^-Cre-ER^T2^: GGGGCCACTAGGGACAGGAT

The H3.1-iCOUNT mouse homology plasmid was generated fusing: 405nt upstream of STOP codon of H3c6, LoxP (5’-ATAACTTCGTATAGCATACATTATACGAAGTTAT-3’), mCherry (without STOP codon; derived from pmCherry-N1), T2A (5’-GGCAGCGGAGAGGGCAGAGGATCCCTCCTCACCTGCGGCGACGTCGAGGAGAACCCCGGACCG-3’), Neo/Kan antibiotic resistance (with STOP codon and 3’UTR, derived from pmCherry-N1), loxP, eGFP (with STOP codon and 3’UTR, derived from pEGFP-N1) and 1kb downstream and integrating it into the pFA6 (pFA6-hphNT1, Euroscarf) backbone cut with HindIII and SacII (NEB) using Gibson Assembly (NEB). The NUP155-iCOUNT homology plasmid was generated by exchanging the homology arms of H3.1-iCOUNT with 1kb upstream of STOP codon and 1kb downstream including STOP codon. The miCOUNT cassette was generated by first fusing: FRT (GAAGTTCCTATTCTCTAGAAAGTATAGGAACTTC), LoxP, tdTomato (without STOP codon), T2A, Neo/Kan antibiotic resistance (with STOP codon and 3’UTR), LoxP, mNeonGreen (with STOP codon and 3’UTR; derived from pNCS-mNeonGreen, Allele Biotech), FRT, Cerulean (with STOP codon and 3’UTR; derived from Cerulean-N1), then adding 405nt upstream of STOP codon and 1kb downstream including STOP codon, all in the pFA6 backbone as described above. The human H3.1-miCOUNT was generated by exchanging the homology arms of the mouse H3.1-miCOUNT with the human homology sequences. Human NUP155-miCOUNT was generated using a synthesized peptide containing: a flexible linker (GGTGGTGGCGGTTCAGGCGGAGGTGGCTCTGGCGGTGGCGGATCG), FRT, LoxP, tdTomato (without STOP codon), 3 Flag tags, T2A, Neo/Kan antibiotic resistance (including STOP codon and 3’UTR), LoxP, eGFP (including STOP codon and 3’UTR), FRT, tagBFP and a V5 tag (including STOP codon and 3’UTR). 1kb upstream and downstream of STOP codon of NUP155 was added and assembled in pFA6 using Gibson Assembly. The PAM sites were mutated in all homology constructs to prevent cutting. To generate the ER^T2^-Cre-ER^T2^ expressing hESC line, DR-GFP from pAAVS1-DRGFP (Addgene No 113193) was replaced by ER^T2^-Cre-ER^T2^ (derived from Addgene No 13777).

To integrate the cassettes, mESCs were co-transfected and mNSPCs/hESCs were co-electroporated with the gRNA plasmid (simultaneously expressing Cas9) and the corresponding homology plasmid. ESCs were selected for correct integration of the iCOUNT cassette using G418 sulfate (300μg/ml for mESCs and 100μg/ml for mNSPCs and hESCs; Gibco) or Puromycin Dihydrochloride (1μg/ml; Gibco) for integration of ER^T2^-Cre-ER^T2^ respectively, until resistant colonies were visible and all cells were dead on a control plate. Correct integration was verified using PCR and fluorescence imaging. The following PCRs were used to check correct integration and to distinguish between *wild-type* (wt), heterozygous and homozygous clones (or animals, see below):

Mouse H3.1-iCOUNT:
Forward: 5’-CAACCAATCAGGAGCATGTT-3’;
Reverse: 5’-TAGGTAGCAGCACTGGCTTG-3’ (wt: 1016bp, H3.1-iCOUNT: 3850bp)
Forward: 5’-CAACCAATCAGGAGCATGTT-3’;
Reverse: 5’-CCCTTGGTCACCTTCAGCTT-3’(wt: no band, H3.1-iCOUNT: 687bp)
Mouse NUP155-iCOUNT:
Forward: 5’-CTGGAACAGAGTGAAAAGCCC-3’
Reverse: 5’-TCTCACTGTGTAGCCCCTGG-3’ (wt: 1540bp, NUP155-iCOUNT: 4388bp)
Forward: 5’-CTGGAACAGAGTGAAAAGCCC-3’
Reverse: 5’-CCCTTGGTCACCTTCAGCTT-3’ (wt: no band, NUP155-iCOUNT: 1229bp)
Human H3.1-miCOUNT:
Forward: 5’-AAC GCG GTA TTT TTG AAT CG-3’
Reverse: 5’-GAT GAC CTC CTC GCC CTT G-3’ (wt: no band, H3.1-miCOUNT:848bp)
Human NUP155-miCOUNT:
Forward: 5’-GGATCCATTCTGGAACAGAA-3’
Reverse: 5’-GAT GAC CTC CTC GCC CTT G-3’ (wt: no band, NUP155-miCOUNT: 1228bp)

### Generation of H3.1-iCOUNT mice

C57BL/6 mice expressing H3.1-iCOUNT were generated by PolyGene (Switzerland). C57BL/6-derived ESCs were transfected with the iCOUNT cassette targeting H3.1 as described above and selected using G418. Single clones were picked, expanded and correct insertion was verified by PCR. Positive cells were injected into blastocysts and 6 chimeras were born. Crossing three chimeras to grey C57BL/6 females yielded heterozygous H3.1-iCOUNT positive animals. To generate homozygous H3.1-iCOUNT mice, H3.1-iCOUNT +/− mice were crossed with each other. Alternatively, H3.1-iCOUNT animals were crossed to Gli1:Cre-ER^T2^ mice (Jackson Lab No 007913: Ahn and Joyner, 2004) and ROSA26:Cre-ER^T2^ mice (Jackson Lab No 008463; Ventura et al., 2007). Animals were always genotyped to distinguish wt, heterozygous and homozygous H3.1-iCOUNT animals (see PCR above). H3.1 tagging caused mosaic expression of the RITE cassette, as there 9 different genes encoding for the histone-variants H3.1 and H3.2. within the *HIST1* gene cluster on chromosome 13 (Wang et al., 1996). Indeed, we observed mosaic expression and mice that had the RITE cassette correctly inserted (verified via genomic PCR) but did not express the transgene.

### Animal handling

All animal experiments were approved by the veterinary office of the Canton of Zurich, Switzerland. To compare iCOUNT-derived data to intravital imaging data we used Gli1:CreER^T2^/tdTom mice, that were derived by crossing Gli1:CreER^T2^ mice with CAG:tdTomato (Jackson Lab, 007914) reporter mice. Two weeks after the placement of the window sparse labelling of NSCs was achieved by single intraperitoneal injection of Tamoxifen (60-70mg/kg bodyweight; Sigma-Aldrich). Imaging followed previously described protocols (Pilz et al., 2018). All mice were kept in individually ventilated cages together with littermates under a 12h light/dark cycle and were provided with water and food *ad libitum*.

### Fluorescence live cell imaging

All images were acquired using a confocal microscope (LSM 800, Zeiss) equipped with an incubator box to maintain 37°C and 5% CO_2_. Prior to imaging, mESCs were transfected with Cre expressing plasmids (pCAG-Cre, Addgene No 13775) and were plated 6h later on Lab Tek II chambered coverglasses (Thermo Fisher Scientific) coated with Poly-L-ornithine (10μg/ml, Sigma-Aldrich) and Laminin (5μg/ml, Sigma-Aldrich). mESCs were imaged every 20min for 64h (H3.1-iCOUNT) or for 38h (NUP155-iCOUNT). mESCs expressing H3.1-miCOUNT were transfected with Cre and Flp-ER^T2^ (pCAG-FlpeERT2, Addgene No 14756) simultaneously, plated after 6h, imaged every 40min for 24h before media was exchanged to TAM (0.5μM 4-hydroxytamoxifen, Sigma-Aldrich) containing media and imaging continued for 44h. hESCs were plated on Matrigel coated chambered coverglasses immediately after electroporation with Cre and 6h later were imaged in standard media every 30min for 70h. Media was exchanged every 24h. Organoids were injected with Cre (1μg/μl) using a FemtoJet 4i microinjector (Eppendorf) and electroporated using the AMAXA Nucleofector device. Organoids were embedded into Phenol Red-free, growth factor reduced Matrigel (Corning) on chambered coverglasses. After polymerization of the Matrigel at 37°C for 30min, imaging media was added that comprised of “Forebrain third medium” (Qian et al., 2016) with DMEM:F12 being exchanged out for Phenol Red-free DMEM:F12 (Thermo Fisher Scientific). Images were taken every 10min for 56h and media was exchanged every 24h.

### Live cell imaging analysis

Movies were analysed using Fiji. Lineages of four starting cells were tracked throughout the movie. Whole nuclear fluorescence (integrated density) was measured and background was subtracted for every time point. Raw values were aligned by the first time point after cell division and by appearance of green fluorescence. Next, red and green fluorescence were normalized to the highest value measured throughout the experiment within one cell lineage and mean values (± standard deviation, n ≥ 4 cells) were calculated from all time points within one cell division. Theoretical values were plotted using the cell cycle length from the experimental data. The theoretical values for red fluorescence was set to 100 prior to the first division and dropped by half with every subsequent division (100, 50, 25 etc.) whereas the green fluorescence first increased to complete the pool of histones (from 0 to 50), dropped by half after cell division (25), increased again (to 75), dropped by half (37.5) and increased again (87.5) etc. Mean values were calculated accordingly (e.g. 0 to 50 results in a mean value of 25). GFP over total fluorescence was calculated for every time point and plotted against time starting at 0h after every cell division. Cells were grouped by the appearance of green signal leading to groups of 0-1 division, 1-2 divisions etc. The GFP over total fluorescence was then grouped into 0 to 50% (0-1 division), 50% to 75% (1-2 divisions), 75% to 87.5% (2-3 divisions) 87.5% to 93.75% (3-4 divisions) and more than 93.75 (4+ divisions). The iCOUNT-based calculated division numbers were then compared to the actual division numbers.

For the miCOUNT live imaging data, fluorescence of tdTomato, mNeonGreen and Cerulean was measured at every time point and data points of three independent lineage trees were normalized by the max of each fluorescence observed. Data points were then pooled by the first cell division mNeonGreen or Cerulean was observed, respectively, and averages (± standard deviation, n ≥ 3 cells) were plotted.

hESCs were tracked as long as possible, red and green fluorescence was measured and aligned by the time of division, normalized by the max fluorescence intensity observed and plotted over time (± standard deviation, n ≥ 3 cells).

Cells depicting green fluorescence were tracked within the organoid as long as possible. Green and red fluorescence of 11 cells was measured for every time point, normalized by the max fluorescence and then plotted over time (± standard deviation, n = 11 cells). Measurements were aligned by the time of cell division. To assess the symmetric distribution of red and green fluorescence between daughter cells, the ratio of green over red fluorescence was measured in daughter cells right after division and values were pair-wise plotted (n ≥ 9 cells).

### Immunostaining

For the embryonic cell division analyses, time-mated H3.1-iCOUNTxROSA26:CreER^T2^ mice received a single intraperitoneal injection of tamoxifen (180mg/kg, Sigma-Aldrich, dissolved in corn oil, Sigma-Aldrich) at different embryonic time points as indicated in figures. Whole embryos (at E11.5 and E15) and embryonic brains (at E15.5, E16, E16.5 and E19) were dissected and fixed overnight in 4% PFA at 4°C. They were first cryoprotected in 15% sucrose at 4°C (overnight) and then in 30% sucrose at 4°C (overnight). Tissues were frozen in OCT compound (Tissue-Tek, Sakura) and stored at −80°C until they were serially sectioned at 20μm using a cryostat (Microm). The slides with frozen sections were stained using methods as described previously (Wegleiter et al., 2019). Tissues were stained against mCherry (1:250, rabbit, Thermo Fisher Scientific), GFP (1:250, goat, Rockland), KI67 (1:250, rat, eBioscience), SOX2 (1:200, rat, eBioscience), SOX2 (1:200, goat, Santa Cruz), TBR2 (1:200, rat, eBioscience), TBR2 (1:200, rabbit, Abcam), CTIP2 (1:200, rat, Abcam) and nuclei were counterstained using 4’,6-Diamidine-2’-phenylindole dihydrochloride (DAPI, 1:5000; Sigma-Aldrich). To assess the effects of staining, mESCs were transfected with Cre, cells were fixed 48h post transfection using 4% PFA for 5min and were stained using the same antibodies against mCherry and GFP. To investigate adult tissues, animals received a lethal dose of Esconarkon (Streuli) followed by trans-cardial perfusion using first 0.9% saline and then 4% PFA in phosphate buffer (0.1 M phosphate, Sigma-Aldrich). Tissues were collected and post-fixed overnight in 4% PFA at 4°C, followed by two nights in 30% Sucrose. Tissues were frozen, cut into 40μm sections using a cryotome (Leica) and stained as described previously (Wegleiter et al., 2019) using the same antibodies against mCherry and GFP. Nuclei were counterstained using DAPI (1:1000).

### Multiplexed immunostaining

To assess the cell division history of adult neural stem cells, adult H3.1-iCOUNTxGli1:CreER^T2^ mice were injected with TAM (180mg/kg) three times constitutively (with one day intervals) and animals were sacrificed by perfusion two weeks after the first TAM injection. Brains were sectioned coronally into 40μm sections that were mounted on poly-D-lysine (PDL, Sigma-Aldrich) pre-treated glass-bottomed 24 well-plates (Cellvis). Sections were stained using iterative indirect immunofluorescence imaging (4i) as described previously (Gut et al., 2018), adapted for fresh-frozen tissue. Tissues were stained for mCherry (1:250, rabbit, Thermo Fisher Scientific), GFP (1:250, goat, Rockland), HOPX (1:250, mouse, Santa Cruz), GFAP (1:500, chicken, Novus), SOX2 (1:250, rat, Invitrogen), S100β (1:500, rabbit, Abcam), DCX (1:350, goat, Santa Cruz), KI67 (1:250, rat, eBioscience), OLIG2 (1:500, rabbit, Millipore), IBA1 (1:500, rabbit, Wako) and NEUN (1:250, mouse, Millipore). During imaging, fluorescent cross-linking was prevented by using phosphate buffer containing N-Acetyl-Cysteine (0.7M, Sigma-Aldrich) at pH 7.4. After imaging, antibodies were eluted using a buffer (pH 2.5) containing L-Glycine (0.5M, Sigma-Aldrich), Urea (3M, Sigma-Aldrich), Guanidinium Chloride (3M, Sigma-Aldrich) and TCEP-HCL (0.07M, Sigma-Aldrich). Tissues were washed and next round of antibodies was applied using a standard staining protocol (Wegleiter et al., 2019).

To stain different cell types in human organoids, day 32 old organoids were injected with Cre plasmid (1μg/μl) and electroporated using the AMAXA nucleofector device. 7 days post Cre injection, organoids were fixed using 4% PFA for 30min, transferred to 30% sucrose solution, embedded in OCT compound and frozen using liquid nitrogen. Organoids were sectioned into 40μm slices using a cryostat, collected in PBS and then mounted on PDL coated glass-bottomed 24 well-plates as described above. 4i was performed using antibodies against SOX2 (1:100, mouse, R&D Systems), KI67 (1:100, mouse, BD-Pharmingen), TBR2 (1:100, rabbit, Abcam), PAX6 (1:100, rabbit, Biolegend), DCX (1:300, goat, Santa Cruz), CTIP2 (1:100, rat, Abcam) and TUJ1 (1:150, mouse, Biolegend). Nuclei were counterstained using DAPI (1:1000).

### Fixed cell image analysis

Images were analysed using Fiji. 4i images of different rounds of staining were aligned by DAPI signal and assembled using Fiji. Whole nuclear fluorescence (integrated density) was measured and background was subtracted for every cell independently. Red fluorescence was normalized by the average of 10 darkest (set to 0%) and 10 brightest “red-only” nuclei (set to 100%) in the analysed tissue. Green fluorescence was normalized by the brightest value (set to 100%) in the analysed tissue. Percentage green fluorescence was calculated by dividing the green fluorescence by total fluorescence (sum of green and red fluorescence). Percentage of green fluorescence was calculated for every cell and plotted grouped by different cell type or localization as indicated. Predicted division number was calculated depending on the percentage green fluorescence ranging from 0 to 50% (0-1 division), 50% to 75% (1-2 divisions), 75% to 87.5% (2-3 divisions) 87.5% to 93.75% (3-4 divisions) and more than 93.75 (4+ divisions). Values of single cells (± standard deviation, n ≥ 28 cells (Figure 3D), n ≥ 21 cells (Figure 4B), n ≥ 90 cells (Figure 4E), n ≥ 10 cells (Figure 5E), n ≥ 15 cells (Figure 6E) were plotted derived from ≥ 3 mice, embryos or organoids.

To assess the symmetric distribution of red and green fluorescence between daughter cells, the ratio of green over red fluorescence was measured on both sides of anaphase cells and values were pair-wise plotted (n = 7 cells). To test the effect of staining, the percentage of green fluorescence was measured in fixed mESCs stained against GFP (using 405nm secondary antibodies) and mCherry (using Cy5 secondary antibodies, Jackson Immuno Research). The percentage of green fluorescence from the endogenous signal (GFP/mCherry) was pair-wise plotted to the percentage of green fluorescence from stainings (405 nm/Cy5, n = 13 cells).

### Theory – cell division histories during cortical development

To assess the utility of the iCOUNT system as a quantitative tool to study cell cycle progression, we made use of existing knowledge of the clonal dynamics during mouse cortical neurogenesis to examine the distribution of cell cycle number. To prepare the analysis, we first considered the predicted distribution using the results of quantitative clonal fate studies by Gao and colleagues, based on the MADM reporter system (*12*) (for details, see Supplemental Computational Modeling). The iCOUNT system predicts that after n rounds of cell division, the mCherry fluorescence intensity signal is a factor of 1/2^n^ smaller than its original level as it becomes replaced by GFP signal. From the ratio of the fluorescence intensity of the mCherry and GFP, we obtained an estimate of the cell division number. Therefore, defining 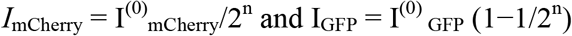, we have the ratio

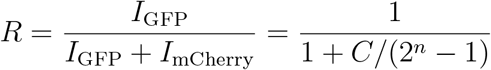

where 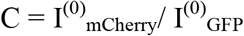. If the baseline flourescence signals are equally intense for both reporters, *C* = 1, and n = − ln(1 − *R*) / ln2. Based on this analysis, the predicted distribution of cell cycle number from the iCOUNT data from the E11.5-E19.5 chase and from the E14.5-E16.5 chase was compared with theoretical predictions from a model obtained from the clone fate analysis of Gao et al. Further details on the comparison are described in the Supplemental Computational Modeling.

### Statistical Analysis

All statistical analyses were done using Prism 8 (GraphPad). First, data was analysed using a D’Agostino & Pearson normality test. If data was normally distributed, an ordinary One-way ANOVA was performed followed by Tukey’s multiple comparisons. If not, a Kruskal-Wallis test was performed followed by Dunn’s multiple comparisons. To compare only two groups, a two-tailed Mann-Whitney test was used. To investigate the symmetric distribution of histones after division, a paired two-tailed t-Test was used. To compare the distribution of R, NR and neurons from iCOUNT and *in vivo* imaging, a Kolmogorov-Smirnov test was used. No samples were measured repeatedly. Figures 1D, 1E and S1C as well as S6A and S6B are different quantifications derived from the same measurements. For all experiments, animals were randomly selected and experimenters were blinded for analysis whenever possible.

### FACS analysis of in vitro samples

mESCs expressing H3.1-iCOUNT or NUP155-iCOUNT were transfected with Cre, whereas mNSPCs expressing H3.1-iCOUNT were electroporated with Cre, plated on Laminin coated plates, re-plated the next day and 6h later media was exchanged to keep proliferation, induce differentiation or quiescence (as described above). At the time points indicated on the figures, mNSPCs and mESCs were collected using TriplE (Thermo Fisher Scientific). Cells were stained with the live/dead marker Zombie Violet (Biolegend) and analysed on an LSR II Fortessa (BD Biosciences). GFP and mCherry expression levels were quantified in singlet-gated live cells using FlowJo (Tree Star). hESCs expressing NUP155-miCOUNT ER^T2^-Cre-ER^T2^ were first treated with TAM (2.5μM 4-hydroxytamoxifen, Sigma) for two days before electroporation with Flp (pCAG-Flpe, Addgene No 13787). Two days post electroporation, cells were collected using Accutase, stained with Zombie NIR (Biolegend) and analysed using the LSR II Fortessa. GFP, tdTom and BFP expression levels were quantified in singlet-gated live cells using FCS Express.

### FACS analysis of mouse tissues

The nuclei dissociation protocol was similar to the previously described protocol (Jaeger et al., 2018). Briefly, the dentate gyrus was carefully excised and immediately placed into a nuclei isolation medium (0.25M sucrose, 25mM KCl, 5mM MgCl2, 10mM TrisHCl, 100mM dithiothreitol, 0.1% Triton, protease inhibitors). Tissue was Dounce homogenized, allowing for mechanical separation of nuclei from cells. The nucleic acid stain Hoechst 33342 (5μM, Life Technologies) was included in the media to facilitate visualization of the nuclei for quantification. Samples were washed, resuspended in nuclei storage buffer (0.167M sucrose, 5mM MgCl2, 10mM TrisHCl, 100mM dithiothreitol, protease inhibitors) and filtered. Solutions and samples were kept cold throughout the protocol. For RNA-seq experiments, tools and solutions were made RNAse-free and RNAse inhibitors were used (Promega, 1:1000 in both isolation and storage buffers). Bone marrow cells were flushed from the tibiae and femurs of mice and red blood cells were lysed using the RBC Lysis Buffer (Biolegend). Hoechst 33342 was added for live cell discrimination. Skin cells were isolated using a modified version of the protocol by Kostic and colleagues (Kostic et al., 2017). Briefly, the dorsal skin of mice was dissected and the connective tissues and fat were removed. The skin was then incubated in PBS without Ca^2+^ and Mg^2+^ (DPBS^--^) with 0.25% Trypsin at 37°C. 2h later, the hair was scrapped off and the skin was triturated using 10 ml pipettes. The cell suspension was filtered successively through 70 and 40 μm cell strainers (Sigma-Aldrich) and washed twice with DPBS^--^ with 3% chelated FBS. Hoechst 33342 was added for live cell discrimination. Dentate gyrus, bone marrow and skin samples were analysed on an LSR II Fortessa (BD Biosciences). GFP and mCherry expression levels were quantified in singlet-gated live cells using FlowJo (Tree Star).

### Single-cell and single-nuclei sorting

Tamoxifen (180mg/kg) was injected into pregnant H3.1-iCOUNT x ROSA26:CreER^T2^ mice at E13.5. 38h later, embryonic cortices were dissected and the nuclei were dissociated as described above (see FACS analysis of mouse tissues). Organoids were injected with Cre (1μg/μl, as described above) and electroporated using the AMAXA Nucleofector device. 4 and 7 days after electroporation (39 days in culture), cells were dissociated as described before (Camp et al., 2015). Short, cells were dissociated using Accutase (Sigma-Aldrich) supplemented with DNAse at 37°C for 25min with gentle mixing every 5min. Accutase was removed by washing and cells were resuspended in DPBS containing EDTA (1mM, Sigma-Aldrich) prior to sorting. Singlet-gated live cells were sorted into 384 well plates containing lysis buffer using a FACSAria III sorter (BD Biosciences). Sorted plates were stored at −80°C until library preparation.

### Single-cell library preparation and sequencing

Library preparation was performed using a mosquito robot HV genomics (TTP Labtech Ltd) following the Smart-seq2 protocol (Picelli et al., 2014). Briefly, 384 well plates containing sorted single nuclei in lysis buffer were thawed and reverse transcription with Superscript II (Thermo Fisher Scientific) and PCR using KAPA Hifi HotStart ReadyMix (Kapa) were performed with the following biotinylated primers (Qiagen): Oligodt (AA GCA GTG GTA TCA ACG CAG AGT ACT TTT TTT TTT TTT TTT TTT TTT TTT TTT TTV N), TSO (AAG CAG TGG TAT CAA CGC AGA GTA CATr GrG+G) and ISPCR primers (AA GCA GTG GTA TCAACG CAG AGT). Following RT-PCR, clean up with Agencourt AMPure XP beads (Beckman Coulter) was carried out and sample concentrations were measured using Bioanalyzer (Agilent Technologies) and normalized at a concentration of 0.3 ng/μl. The Nextera XT DNA library prep kit (Illumina) was used for subsequent sample preparation. Samples were subjected to a tagmentation reaction, indexing, and PCR amplified. Libraries were then mixed in 384-sample pools and purified with Agencourt AMPure XP beads. Ready DNA libraries were quality controlled using D1000 Screen Tape Assay (Agilent Technologies). Samples were sequenced at the Functional Genomics Center Zurich on Illumina HiSeq 2500 or HiSeq4000 sequencers with single-end 125bp reads.

### Single-cell RNA-seq analysis

Single-end 126nt-long reads were adaptor removed and trimmed using cutadapt v1.16 and sickle v1.33 with default parameters. Reads were mapped against the mouse GRCm38.90 primary assembly or the human GRCh38.p13 primary assembly using STAR v2.6.0c (Dobin et al., 2013) in `alignReads` mode. Gene count matrices were quantified at the exon level ignoring multimappers using featurecounts from subread v1.6.2. RNA velocity loom files were generated with velocyto v0.17.17 (La Manno et al., 2018) in `run-smartseq2` mode.

Nuclei from the mouse embryonic cortex were quality checked using scater v1.12.2 (McCarthy et al., 2017). Only cells with 1000-3000 genes detected, less than 5% of mitochondrial reads and more than 50’000 reads were kept. In each dataset (i.e. 384 well-plate), nuclei with more or less than 1.5-fold the median read content per cell were excluded. A total of 552 nuclei passed QC filtering. 17 interneuron nuclei (*Gad2^+^Gad1*^+^) were filtered out. A total of 535 nuclei were retained for final analysis. Human organoid cells were quality checked using scater v1.12.2 (McCarthy et al., 2017). Only cells with 3000-7000 genes detected, less than 10% of mitochondrial reads and more than 50’000 reads were kept. In each dataset (i.e. 384 well-plate), cells with more or less than 1.5-fold the median read content per cell were excluded. A total of 641 cells passed QC filtering. 207 RSPO cells (*RSPO2^+^*) were filtered out. A total of 434 cells were retained for final analysis. Seurat v3.0.2 was used to normalize read counts, regress out the library size and mitochondrial reads proportion per cell, find variable features with the `vst` method, integrate multiple plates as in (Stuart et al., 2019), dimensionality reduce the data, cluster cells and test for differentially expressed genes. Differential gene expression between orange and green NSPCs was tested only for genes detected in at least 20% of the cells in either of the populations. Logistic regression fit with batch as latent variable (null model) and cell type as predictor were run using Seurat's `FindMarkers`. Genes with p-value below 0.05 were deemed significant. Functional enrichment categorization was performed using DAVID 6.8 (Huang da et al., 2009). The GONetwork R package (https://github.com/saralinker/GONetwork) was used to identify the functional categories represented in genes up-regulated in orange or green human organoid NSPCs. Functional categories found in both human organoid cells and mouse embryonic cortex nuclei were highlighted.

## Data availability

All data are available from the authors at request. scRNA-seq data have been submitted into GEO.

## Supplemental Figures

**Figure S1:**
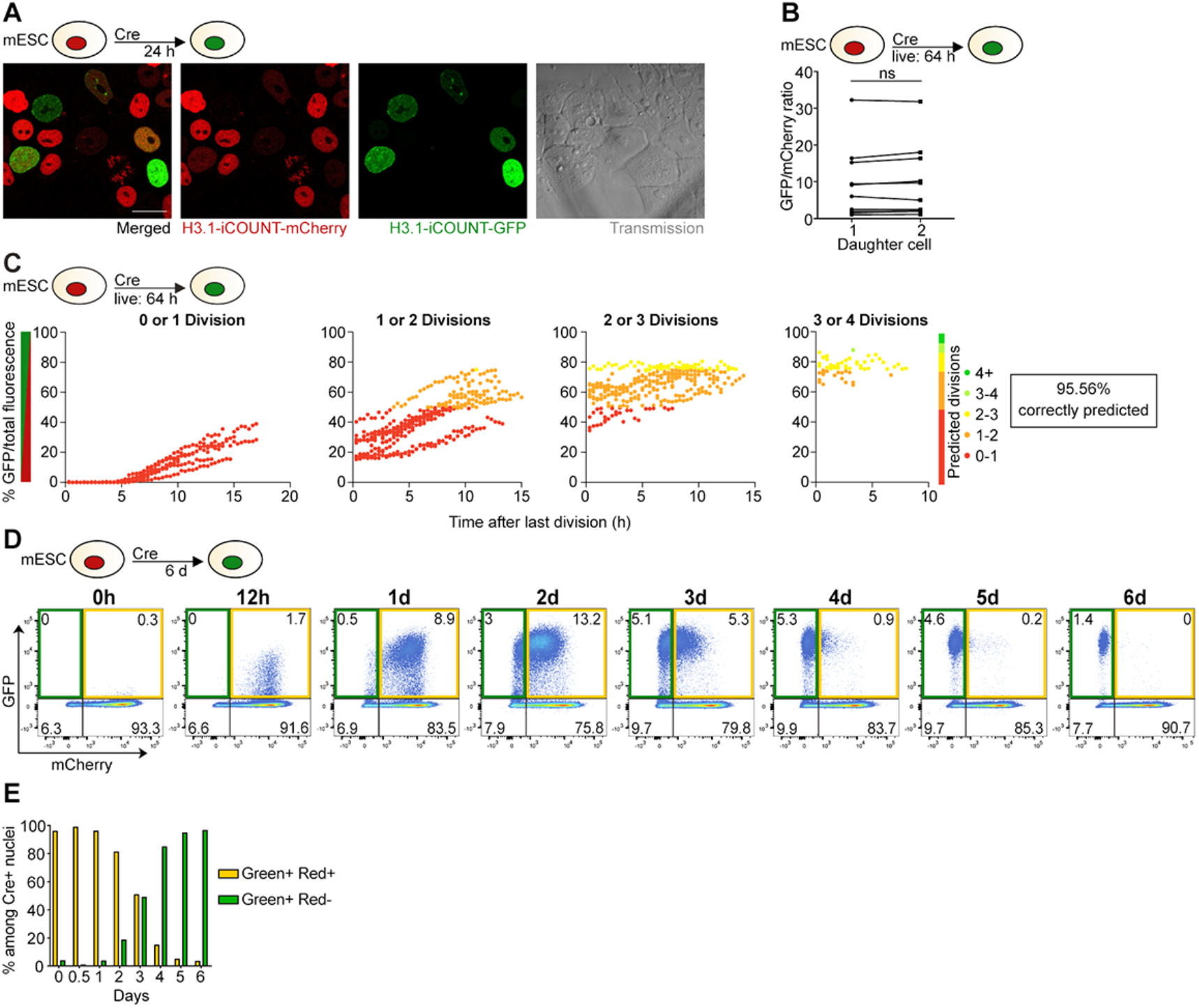
iCOUNT reports cell division events – Related to Figure 1. (A) Expression of mCherry-(red) and GFP-tagged (green) histones in mouse ESCs 24h after Cre-mediated recombination. Left panel shows merged signal. (B) Quantifications show ratios of green over red fluorescent intensities of sister cell pairs. (C) Live cell imaging of iCOUNT mouse ESCs for 64h confirms >95% correct prediction of cell divisions based on fluorescent intensities. Shown are measured data points and their predicted division number for individual observed division bins. (D) FACS analyses show relative shift from red/green to green fluorescent intensities in mouse ESCs in the course of 6d after Cre-mediated recombination. (E) Quantifications of the boxed areas show the gradual change of iCOUNT colour. ns, not significant. Scale bar represents 20μm.

**Figure S2:**
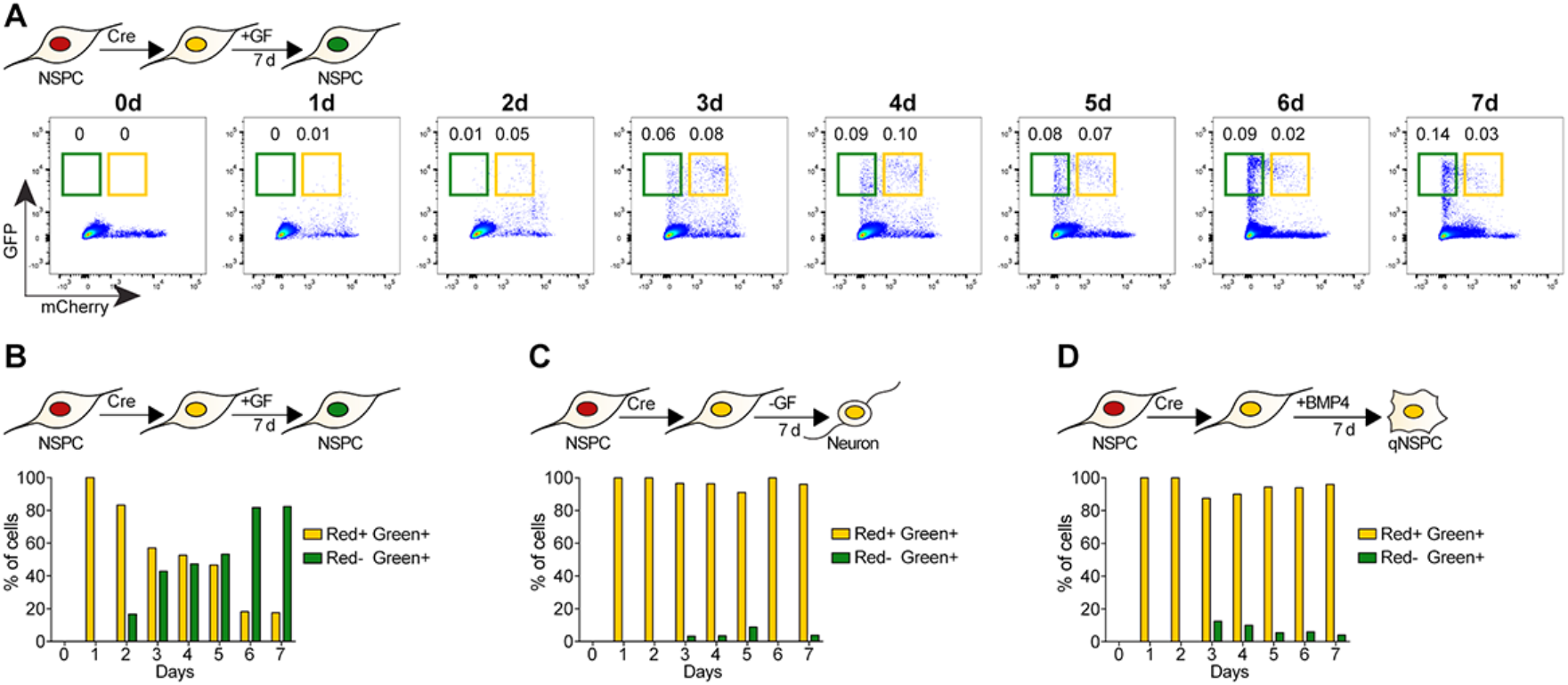
iCOUNT colour exchange in different cell types and conditions – Related to Figure 2. (A) FACS analyses show relative shift from red/green to green fluorescent intensities in mouse NSPCs in the course of 7d after Cre-mediated recombination when exposed to growth factors (GF). (B) Quantification of the boxed areas. (C) In contrast, differentiation by withdrawal of GF (-GF) prevents a fluorescence shift throughout 7d. (D) Inducing quiescence with BMP4 for 7d also prevents a fluorescence shift.

**Figure S3:**
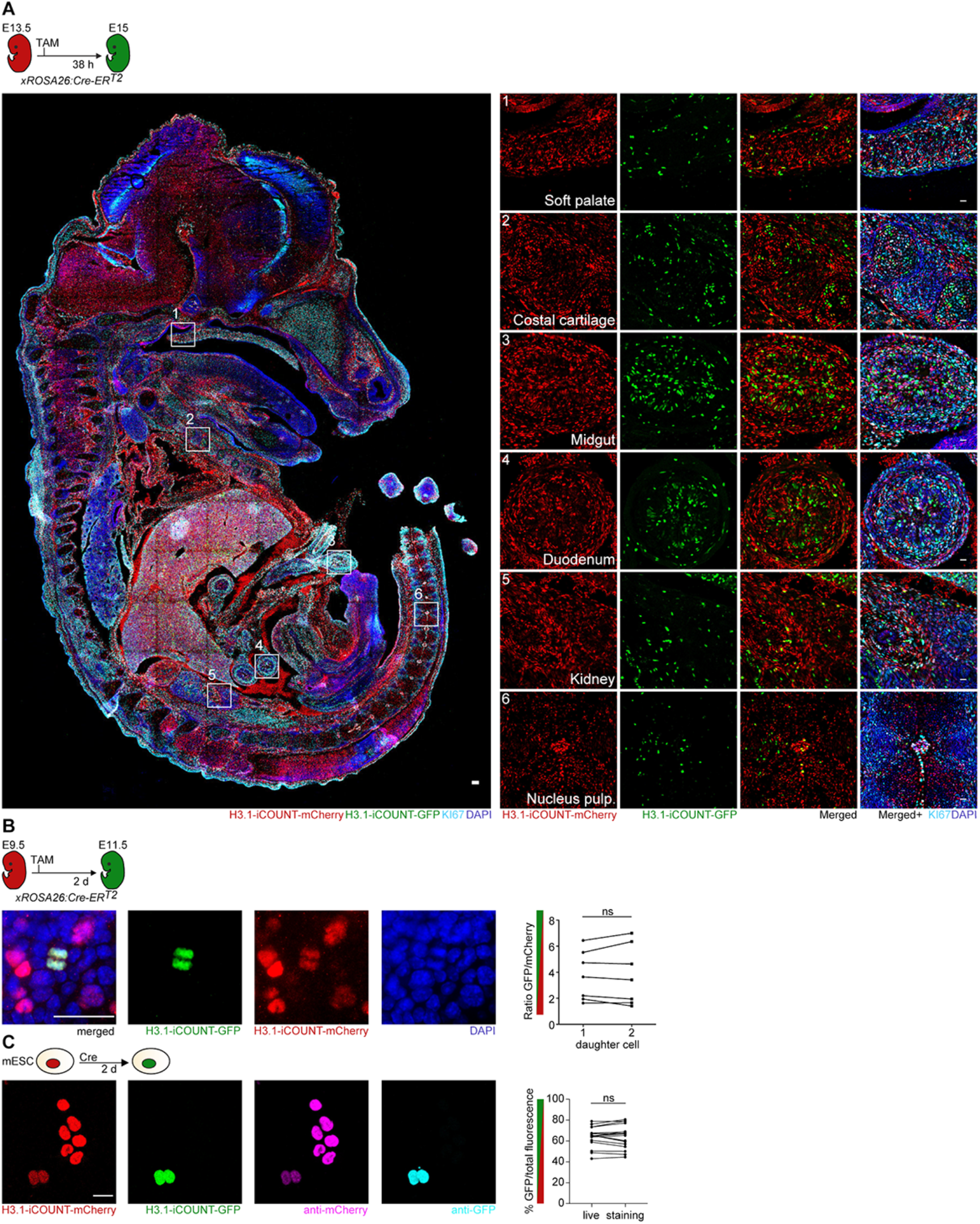
iCOUNT reports cell division history in embryonic tissues – Related to Figure 3. (A) Overview of iCOUNT mouse at E15 (recombination using TAM at E13.5). Right panels show mCherry (red), GFP (green), KI67 (light blue) signal in tissues indicated. Symmetric segregation of mCherry-(red) and GFP-tagged (green) histones in the developing mouse cortex in dividing progenitors. Right panel shows quantification of green over red fluorescent intensities for sister pairs. (C) Amplification of endogenous iCOUNT signal in mESCs of mCherry (red) and GFP (green) using antibodies against mCherry (pink) and GFP (turquoise) maintains relative fluorescence ratios. Quantifications of analysed cells are shown in right panel. Nuclei were counterstained with DAPI (blue). ns, not significant. Scale bars represent 100μm in A (left panel) and 20μm in A (right panels), B-C.

**Figure S4:**
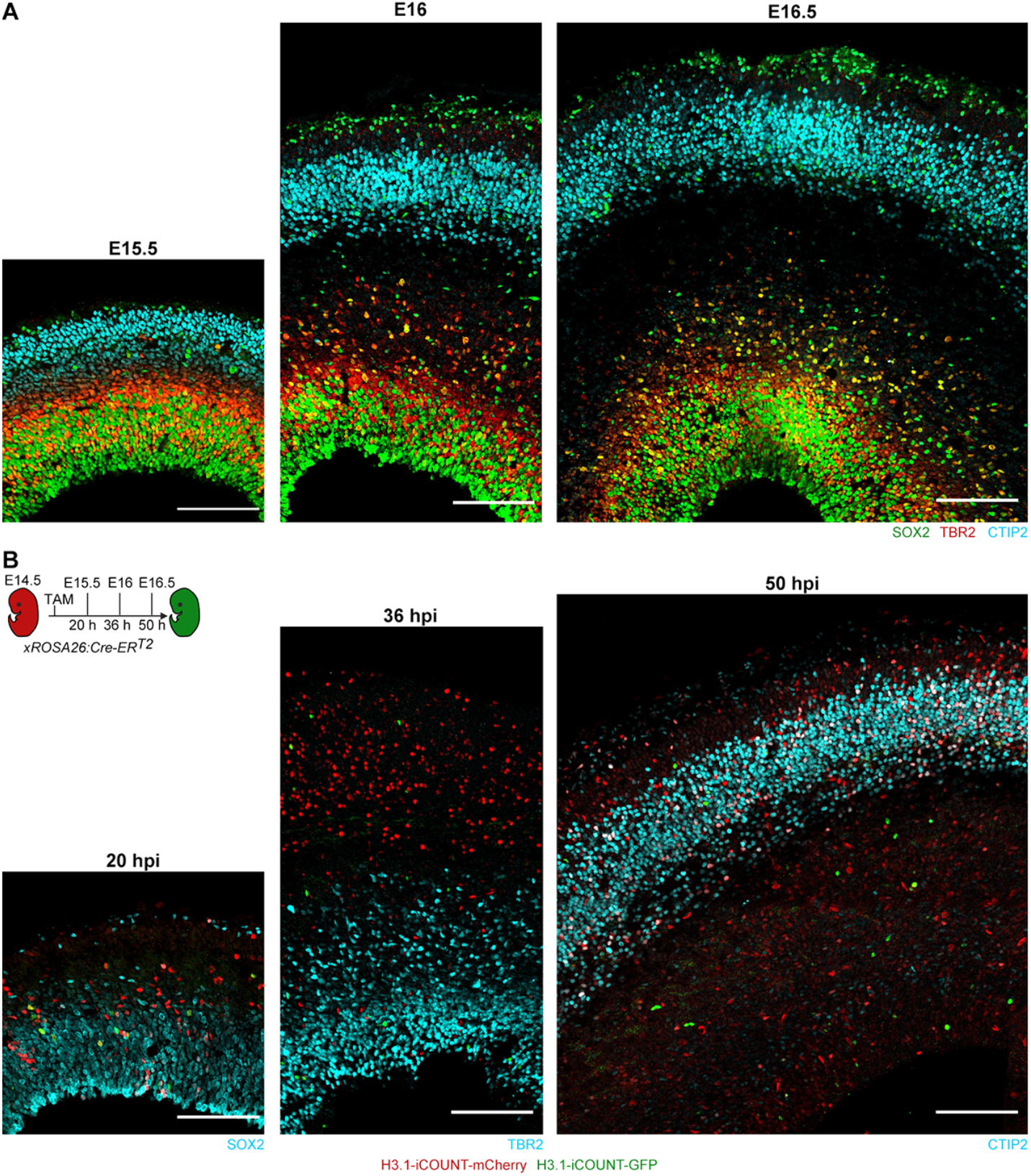
Recombined iCOUNT cells in different areas of the developing cortex – Related to Figure 4. (A) Images of the developing mouse cortex analysed at E15.5, E16 and E16.5 stained with SOX2 (green), TBR2 (red) and CTIP2 (light blue) depicting different layers. (B) Representative images showing iCOUNT-labelled cells containing H3.1-mCherry (red), H3.1-GFP (green) and SOX2, TBR2 or CTIP2 (light blue) in the developing cortex of iCOUNT mice induced at E14.5 and analysed at indicated time points. Scale bars represent 100μm.

**Figure S5:**
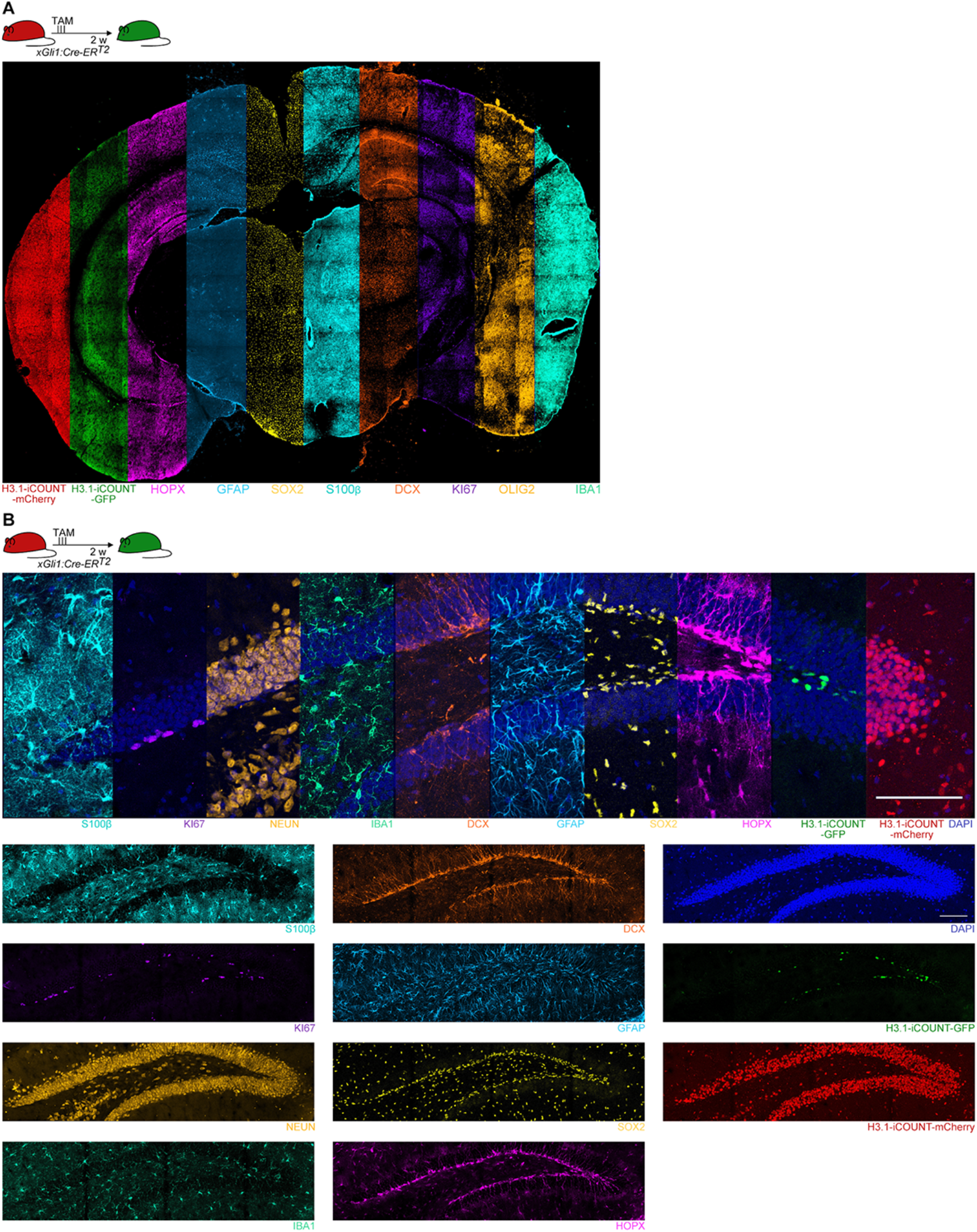
iCOUNT signal in adult mouse tissues – Related to Figure 5. (A) Overview of an adult iCOUNT mouse brain with expression of the tagged H3.1 and analysed using 4i technology with the markers indicated. (B) High power view of the mouse hippocampus and analysed using 4i technology with the markers indicated. Nuclei were counterstained with DAPI (blue). Scale bars represent 100μm.

**Figure S6:**
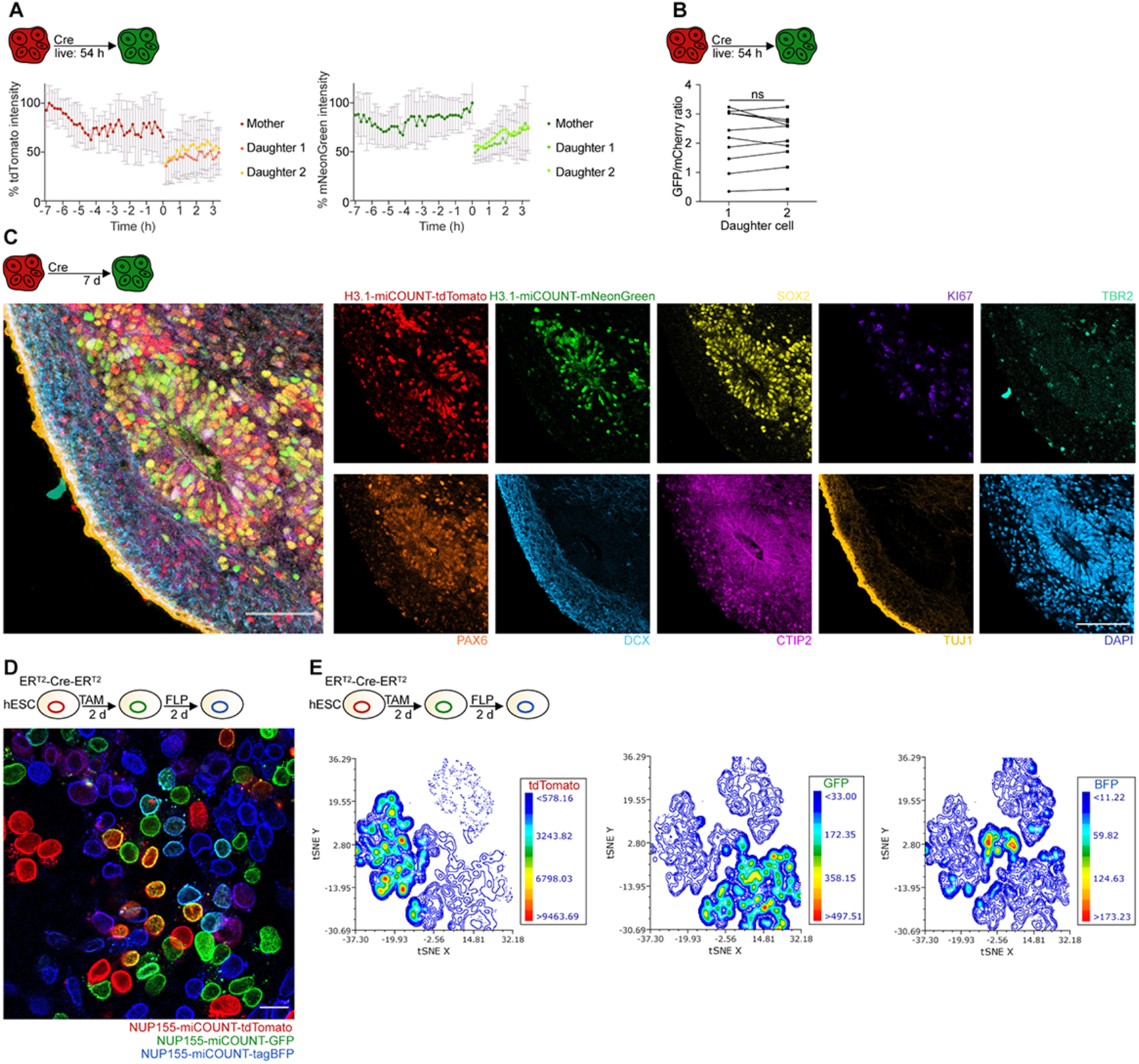
iCOUNT and miCOUNT reports cell division events in human cells – Related to Figure 6. (A) Quantification of measured changes in fluorescence intensities for red and green histones of iCOUNT-targeted cells in the organoid during live imaging aligned by the time of division (mean ± SD). (B) Symmetric segregation of tdTomato-(red) and mNeonGreen-tagged (green) histones in human organoids upon cell division in sister pairs. Graph shows quantification of green over red fluorescent intensities for sister pairs. (C) 4i-based phenotyping of iCOUNT-targeted cells in human ESC-derived forebrain organoids using a panel of protein markers as indicated. (D) NUP155-miCOUNT expressing human ESCs 2d after TAM and 2d after FLIP-based recombination. Note the presence of tdTomato-tagged (red), GFP-tagged (green), and tagBFP-tagged (blue) NUP155 and their combinations in human ESCs. (E) FACS analyses confirm the presence of red, green, blue fluorescent intensities in human NUP155-miCOUNT expressing ESCs, depicted using a tSNE plot of fluorescence values. ns, not significant. Scale bars represent 100μm in C and 20μm in D.

**Figure S7:**
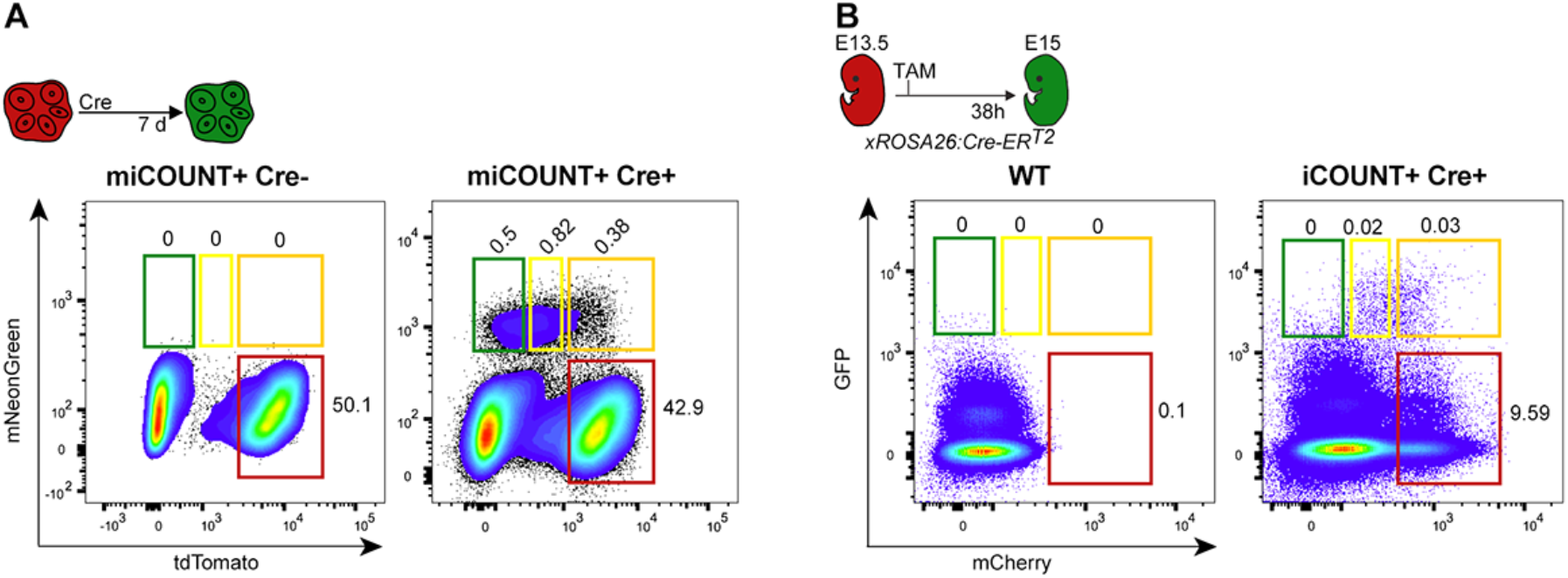
Molecular consequences of previous cell divisions in mouse and human NSPCs – Related to Figure 7. (A) FACS analyses of cells with distinct red/green histone ratios in human forebrain organoids 7d after induction of Cre. Shown are FACS plots of cells without (left) and with induction of Cre (right panel). (B) FACS analyses of cells derived from developing mouse cortex at the time points indicated. Shown are FACS plots of wt control animals (left) and iCOUNT mice 38h after induction of Cre (right panel). Boxed areas in different colours depict gating strategies to collect single cells of different iCOUNT colours used for scRNA-seq.

**Supplemental Computational Modeling – Related to Figure 4.**

## Supplemental Tables and Movies

**Table S1 – Related to Figure 7.** Table of genes up-regulated in green vs. orange human organoid NSPCs.

**Table S2 – Related to Figure 7.** Table of genes up-regulated in orange vs. green human organoid NSPCs.

**Table S3 – Related to Figure 7.** List of GO term clusters generated by genes upregulated in green vs. orange (upper panel) and orange vs. green (lower panel) human organoid NSPCs including the gene names representing each node in the figures (Figure 7E-F).

**Movie S1 – Related to Figure 1.** mESCs expressing H3.1-iCOUNT imaged for 64h starting 6h post transfection with Cre recombinase. Movie shows the gradual change from red (mCherry) to green (GFP)-tagged histones in cells that underwent Cre-dependent recombination. Duration of the movie: 64h. Scale bar represents 100μm.

**Movie S2 – Related to Figure 2.** mESCs expressing H.3-miCOUNT were co-transfected with Cre and TAM-inducible Flp 6h prior to imaging. 24h after imaging started, TAM was added and live imaging continued for additional 44h. The movie shows first a gradual change from red (tdTomato) to green (mNeonGreen) histones followed by a change from green to blue (Cerulean) histones. Duration of the movie: 68h. Scale bar represents 20μm.

**Movie S3 – Related to Figure 6.** Human forebrain-regionalized organoids derived from H3.1-miCOUNT expressing hESCs were imaged for 54h starting 24h post Cre injection. A gradual change from red (tdTomato) to green (mNeonGreen) can be seen in cells that received Cre. Arrow highlights a H3.1-mNeonGreen expressing cell that undergoes interkinetic nuclear migration and divides at the ventricle. Movie is set to slow motion during division. Duration of the movie: 54h. Scale bar represents 20μm.

